# Long-term single-molecule Ca^2+^ flux recordings reveal mode-switching regulation of Ca^2+^-ATPases

**DOI:** 10.64898/2026.01.28.701947

**Authors:** Mads P. Møller, Aleksander Cvjetkovic, Eleftherios Kosmidis, Alessandra Narducci, Michael Isselstein, Christopher G. Shuttle, Bo Justesen, Ida L. Jørgensen, Sara Basse-Hansen, Mateusz Dyla, Jesper Holmkvist, Magnus Kjaergaard, Per Armstrup Pedersen, Poul Nissen, Thomas Günther Pomorski, Dimitrios Stamou

## Abstract

Calcium (Ca²⁺) is a universal second messenger that governs processes ranging from muscle contraction and secretion to gene expression and cell fate. Ca²⁺-ATPases establish and maintain steep Ca²⁺ gradients across intracellular membranes, yet how regulatory inputs modulate the underlying single-pump Ca²⁺ currents has remained inaccessible. Here we develop a non-saturating, self-regenerating single-vesicle assay that monitors over hours the zeptoampere (10⁻²¹ A) currents produced by individual Ca²⁺-ATPases. In parallel, we establish a workflow to record single-molecule currents from human sarco/endoplasmic reticulum Ca²⁺-ATPases (hSERCA) in native endoplasmic reticulum vesicles. Using reconstituted LMCA1, a bacterial SERCA homologue, we observe stochastic switching between minute-long pumping and inactive modes, as well as uncoupled Ca²⁺ leakage events that are suppressed by vanadate. Extravesicular pH controls a previously unrecognized *dormant* pre-activation mode that delays the onset of pumping, without measurably altering pumping rates or active-mode lifetimes. Extending the assay to endogenous hSERCA reveals delayed activation and ultraslow pumping/inactive mode-switching without detectable transprotein Ca²⁺ leakage. ATP and Ca²⁺ regulate the probability of hSERCA activation by modulating dormant-mode occupancy. Together, these results extend ultraslow mode-switching, previously observed only for proton pumps, to Ca²⁺-ATPases and identify probability-gated entry into productive cycling as a distinct regulatory axis of human Ca²⁺-ATPase regulation that can modulate the timing and heterogeneity of Ca²⁺ store refilling without changing on-cycle kinetics.

## Introduction

Primary active transporters convert metabolic energy into transmembrane electrochemical gradients that power diverse biological processes. In contrast to ion channels, individual transporters rarely generate electrical signals accessible to electrophysiology^1^. Mechanistic and regulatory models are therefore largely inferred from ensemble-average measurements of transport rates and from structural snapshots of the transport cycle^2–4^.

Long-duration single-molecule flux recordings recently challenged a core assumption embedded in many ensemble models: that regulation manifests as continuous tuning of catalytic turnover. Instead, two proton pumps were found to stochastically exit their canonical transport cycles into long-lived inactive and leaky *modes* lasting minutes, with regulatory inputs primarily tuning the probability of occupying these modes rather than the intrinsic transport rates within the cycle^5,6^. Long-lived (locked) conformations have also been observed in pioneering single-molecule studies of secondary active transporter homologues^7–10^. However, though conceivably related to modes, such locked states have been interpreted as intermediates of the catalytic cycle, and their connection to physiological regulation is currently unclear.

Proton pumps share specialized transport chemistries and coupling mechanisms that differ from the ones used to transport most other ions and small molecules^11–14^ by harnessing the unique ability of protons to move efficiently through hydrogen-bonded “proton-wiring” networks. It has therefore remained uncertain whether ultraslow mode-switching represents a general regulatory mechanism for transporters or a peculiarity of proton transport. Ca^2+^-ATPases provide a stringent test case. Sarco/endoplasmic reticulum Ca^2+^-ATPases (SERCAs) and their bacterial homologues are archetypal P-type ATPases that follow the classical Post-Albers cycle to pump Ca^2+^ against steep gradients^15–18^. Their activity underlies the essential physiology related to calcium signaling, e.g., muscle relaxation and organellar Ca^2+^ homeostasis, and is controlled by membrane composition, ligands, luminal and cytosolic conditions, and regulatory proteins^19–25^. Single-molecule conformational studies have resolved rapid transitions within the transport cycle over seconds^26,27^. Yet, such measurements did not reveal whether individual Ca^2+^-ATPases also access rare, ultralong-lived off-cycle modes that would only become apparent over many thousands of turnovers.

Here, we introduce a long-duration, non-saturating single-molecule assay that reports the activity of individual Ca^2+^ pumps as ion fluxes into the lumen of single immobilized nanoscopic vesicles. The central challenge is that biological membranes are inherently impermeable to Ca^2+^ (ref. ^28^), which causes pumping to rapidly saturate luminal Ca^2+^ indicators and prevents continuous readout. We overcome this by tuning native membrane Ca^2+^ permeability using the Ca^2+^ ionophore ionomycin such that active pumping drives the vesicle lumen to a dynamic steady state below indicator saturation. At the same time, passive Ca^2+^ efflux resets the signal when the pump becomes inactive. This self-regenerating regime enables uninterrupted recordings of single Ca^2+^-ATPases pumping for hours. Applying this platform to reconstituted LMCA1 and to native hSERCA in native endoplasmic reticulum (ER) vesicles, we uncover ultraslow mode-switching and a previously unrecognized dormant activation mode that underpins regulation of Ca^2+^ pumping. Probabilistic regulation that acts by gating *whether* and *when* pumps enter productive cycling, rather than by continuously tuning the speed of a permanently active cycle, would introduce latency and intermittency in Ca²⁺ sequestration and clearance, and could in principle generate persistent heterogeneity between otherwise similar organelles or cells, even when the intrinsic turnover within the catalytic cycle is unchanged.

### A self-regenerating assay enables long-term single Ca^2+^-ATPase recordings with zepto-ampere resolution

We monitored Ca^2+^ transport by LMCA1 reconstituted (see Methods) in individual nanoscopic vesicles (Fig. S1) tethered to a passivated surface and imaged by fluorescence microscopy^29–31^. Each vesicle encapsulated a Ca^2+^-sensitive indicator, Cal-520 (see also Fig. S2 and Methods), whose intensity reports luminal free Ca^2+^ in the range 0.5 to 100 µM (Fig. 1b). Fluorescence signals were converted to Ca^2+^ using a local single-vesicle calibration method (Fig. S3 - S5 and Supplementary methods). As expected for tightly sealed lipid bilayers, vesicles are effectively impermeable to Ca^2+^, so active pumping, triggered by the addition of ATP, rapidly drives luminal Ca^2+^ beyond the dynamic range of the indicator and irreversibly saturates the signal (Fig. 1c, Fig. S6, Fig. S7 for negative controls, Supplementary discussion).

**Fig. 1.**
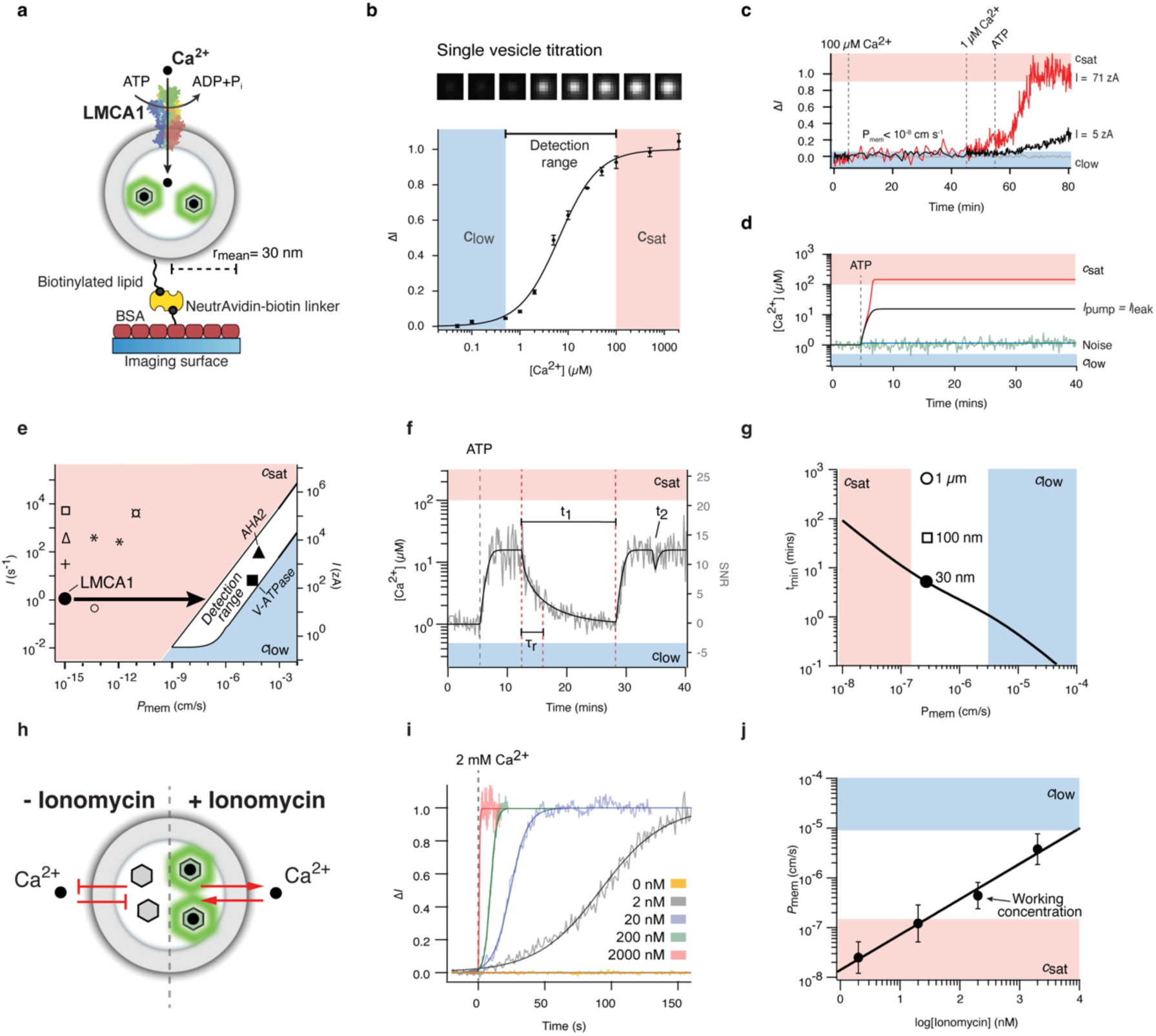
Development of a non-saturating and regenerating Ca-ATPase activity assay with tunable membrane permeability and zepto-ampere resolution. **a,** Illustration of the single-molecule transporter assay using LMCA1-reconstituted vesicles (cytoplasmic side out) with encapsulated fluorescent Ca^2+^ indicators. **b,** Example fluorescent microscopy of single-vesicle dose-response curve of [Ca^2+^] fitted with a sigmoidal function. Vesicles were permeabilized with ionomycin and subjected to different external [Ca^2+^]. Changes in luminal [Ca^2+^] can be accurately monitored over the range 0.5-100 μM (c_low_ and c_sat_, blue and red shaded areas). Error bars correspond to the s.d. of 3 images. **c,** Vesicles are effectively impermeable to Ca^2+^ 1-100 µM Ca^2+^ (∼100-fold) gradient for 35 min. Upon addition of ATP, LMCA1-mediated Ca^2+^ pumping saturated the fluorescent indicator in most vesicles (red trace). Rare ultra-slow Ca^2+^ pumping kinetics that did not reach saturation (black trace). Control trace is shown in grey. **d,** Simulations of fixed Ca^2+^ influx in vesicles with varying Ca^2+^ permeability (*P*_mem_). The plateau reflects the dynamic equilibrium between Ca^2+^ influx and efflux. Changing *P*_mem_ shifts the equilibrium in/out of the sensor detection range. **e,** Simulations reveal that the detection range of the assay scales with *P*_mem_. *P*_mem_ must be increased by ∼9 orders of magnitude to detect Ca^2+^ pumping by LMCA1^32^. The pumping rate and ion substrate permeability of other typical pumps are indicated (+: SPCA1, Δ: SERCA/PMCA, 7: NCX1, O: Glt_Ph_, *: Na^+^/K^+^ pump, ¤: ClC-ec1, ∎: V-ATPase, Δ: AHA2)^5,6,33–37,7,10,38^. **f**, Simulated activity trace with corresponding typical experimental noise displaying mode-switching between active and inactive modes (On, Off) in a Ca^2+^ permeabilized vesicle. Ca^2+^ leaks out of the vesicle (τ_efflux_, see also Supplementary discussion) when the pump switches to an inactive mode, allowing the system to regenerate (t_1_). If the dwell time of the inactive mode is too short, mode-switching is not detected (t_2_). **g**, Simulations reveal the minimum detectable dwell-time range of mode-switching events (t_min_) as a function of *P*_mem_ (black line) for a range of vesicle radii. Mode-switching within these limits will be resolved. **h**, Illustration of ionomycin acting as a Ca²⁺ ionophore to increase membrane permeability. **i**, Representative kinetics of Ca^2+^ influx in single vesicles for different ionomycin concentrations. **j,** Membrane permeability to Ca^2+^ as a function of ionomycin concentration. The white, non-shaded area indicates the range of *P*_mem_ for which a current of ∼300 zA would be resolved. Mean ± s.d. from 400-1000 single vesicle kinetics. *n* = 3 experimental replicates. Hereafter, *n* is the number of replicates.

To establish a sustained readout, we sought conditions in which transporter-mediated Ca^2+^ influx is balanced by passive Ca^2+^ efflux to create a non-equilibrium steady state in the measurable range (Fig. 1d). We used a quantitative physical model of Ca^2+^ accumulation that accounts for pump-driven influx, electrodiffusive leakage, buffering, and the buildup of membrane potential^5,6,39^ (see also Supplementary discussion). Simulations showed that the steady-state luminal Ca^2+^ depends strongly on the membrane Ca^2+^ permeability (*P*_mem_), and that *P*_mem_ must be increased by ∼ 9-10 orders of magnitude relative to a native lipid bilayer to render the zepto-ampere currents of LMCA1^32^ continuously detectable without saturating the indicator (Fig. 1e and Fig. S8).

We therefore used ionomycin, a canonical Ca^2+^ ionophore^40^, to tune *P*_mem_ in a controlled manner (Fig. 1h). By measuring Ca^2+^ influx kinetics in single vesicles at defined ionomycin concentrations and fitting these traces with the physical model, we obtained a calibration of ionomycin concentration to *P*_mem_ (Fig. 1i, j). This calibration, together with simulations of signal-to-noise and response time (see Fig. S9 – S11 and supplementary discussion), defines an experimental window in which the assay is predicted to resolve ∼300 zA Ca^2+^ currents and minute-scale mode-switching events (Fig. 1f, g). The fluorescence response generated by a single-ion translocation event was smaller than the noise level of the kinetic traces (∼60%) and could therefore not be directly resolved as discrete steps in fluorescence intensity (Fig. S8e, f, Supplementary discussion). In the absence of mode switching, extending the recording duration permitted detection of average currents down to a few zA (Fig. S8).

### LMCA1 switches between pumping, inactive, and Ca^2+^-leaky modes

In the tuned-permeability regime, pumping by an individual Ca^2+^-ATPase is expected to increase luminal Ca^2+^ to a steady-state plateau. If the pump becomes inactive, passive Ca^2+^ efflux through ionomycin relaxes the vesicle back toward baseline. The vesicle would thus repeatedly cycle between low- and high-Ca^2+^ modes, making the assay self-regenerating and enabling uninterrupted long-term recordings (Fig. 2a).

**Fig. 2.**
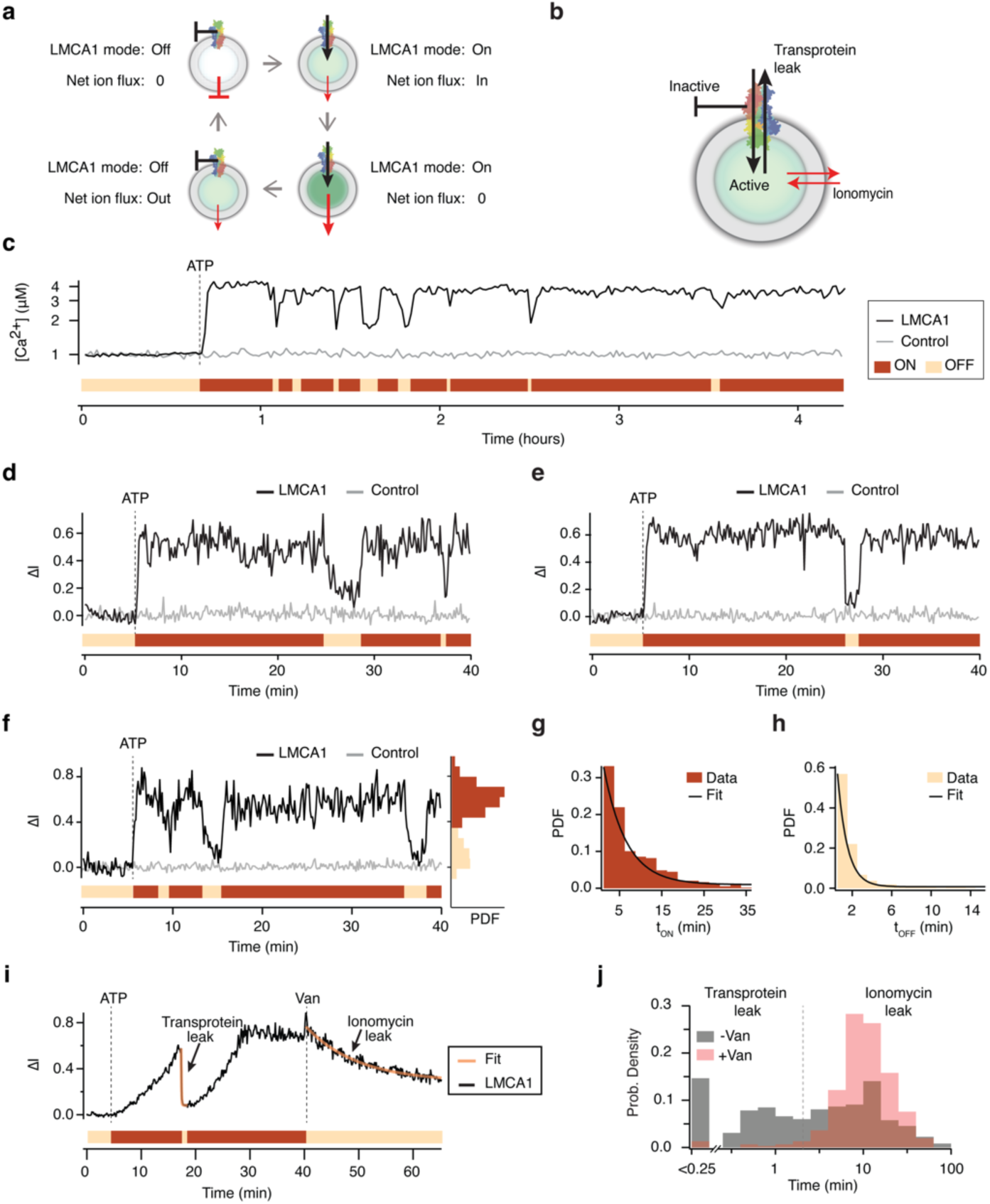
Single-molecule Ca^2+^ transport measurements reveal stochastic switching of LMCA1 between active, inactive, and leaky modes. **a,** Illustration of the self-generating transporter assay for long-term kinetic measurements of Ca^2+^ pumping in the presence of ionomycin. **b,** Illustration depicting a proteoliposome in which a transprotein leakage pathway allows calcium to escape down its concentration gradient through LMCA1. **c,** Example 5-hour calcium pumping kinetics by single LMCA1 molecules displaying ultra-long lived stochastic switching (black line). Example trace of an inactive vesicle (grey line). **d-f**, Zoom-in of kinetic traces showing Ca^2+^ pumping and switching between active and inactive modes by single LMCA1 molecules. In (f) right: histogram of accumulated intensity over time reveals two distinct populations corresponding to active and inactive modes. **g-h**, Histograms of active and inactive mode dwell times fitted with a single exponential function, *1*_ON_ = 326 ± 24 s and *1*_OFF_ = 78 ± 8 s. **i**, Example kinetic trace showcasing a typical transprotein leakage event. The leakage lifetime through the transprotein pathway is an order of magnitude faster than that of ionomycin upon addition of the LMCA1 inhibitor vanadate (Van). **j**, Histogram of leakage lifetimes during LMCA1 activity in the presence of ATP only (grey) and upon addition of Van (red). In the absence of inhibitor leakage lifetime populations distinctly showcase a fast and a slow mechanism of Ca^2+^ efflux. The fast leakage pathway can be attributed to LMCA1 because inhibition of the pump also blocks fast efflux.

Recordings of single LMCA1 molecules in reconstituted proteoliposomes (see Methods and Fig. S12) confirmed stochastic switching between long-lived pumping and inactive modes over multi-hour traces (Fig. 2c-f and Fig. S13, S14), consistent with previous single-molecule pumping measurements of a proton P-type ATPase^5^. Dwell-time analysis showed that both the pumping and inactive modes persist for minutes and are well described by single-exponential distributions with characteristic lifetimes τ_ON_ = 326 ± 24 s and τ_OFF_ = 78 ± 8 s (Fig. 2g, h). These dynamics are ∼1000-fold slower than the full cycle^26^ (∼5 Ca^2+^ s^-1^), suggesting the inactive mode is not part of the cycle. Thus, during our long observation times, LMCA1 appears to exit the Post-Albers transport cycle into an off-cycle inactive state and re-enter productive pumping stochastically.

Beyond pumping and inactivity, we also observed inactive periods in which luminal Ca^2+^ collapsed substantially faster than expected from ionomycin-mediated efflux alone (Fig. 2b, i). Addition of the P-type ATPase inhibitor vanadate^41^ abolished this fast efflux component, leaving only the slower ionomycin-driven leakage (Fig. 2j and Fig. S15). The distribution of efflux lifetimes therefore supports a rare transprotein Ca^2+^ leakage pathway that is suppressed by inhibitor binding. Together, these observations establish that LMCA1 populates multiple ultralong-lived functional modes, including a leaky mode that partially uncouples the pump from strict transport.

### Extravesicular pH regulates LMCA1 activation through a dormant pre-activation mode

LMCA1 is a bacterial P-type Ca^2+^ pump that physiologically promotes alkaline stress tolerance by upregulating Ca^2+^ flux^32^. It shows optimal ATPase activity at alkaline pH^32^. We therefore asked whether extravesicular pH (corresponding to cytosolic pH in the bacterium) regulates LMCA1 by tuning the intrinsic kinetics of Ca^2+^ transport, or instead by controlling access to productive cycling through ultraslow on/off switching. To address this, we monitored luminal Ca^2+^ transport at pH 7.5, followed by a rapid pH jump to 8.5, resulting in a higher ensemble plateau (Fig. 3a). At face value, this could be interpreted as pH accelerating pumping or elevating the steady-state Ca^2+^ level reached by each active transporter. However, single-molecule trajectories did not reveal a statistically significant change in steady-state luminal Ca^2+^ level (Fig. 3b, c). Thus, the increase in the ensemble-average signal upon alkalinization cannot be explained primarily by a pH-driven change in the intrinsic pumping rate or in the steady-state Ca^2+^ level achieved by already active LMCA1 molecules. Similarly, pH did not alter the lifetimes of the inactive/active modes (Fig. 3d).

**Fig. 3.**
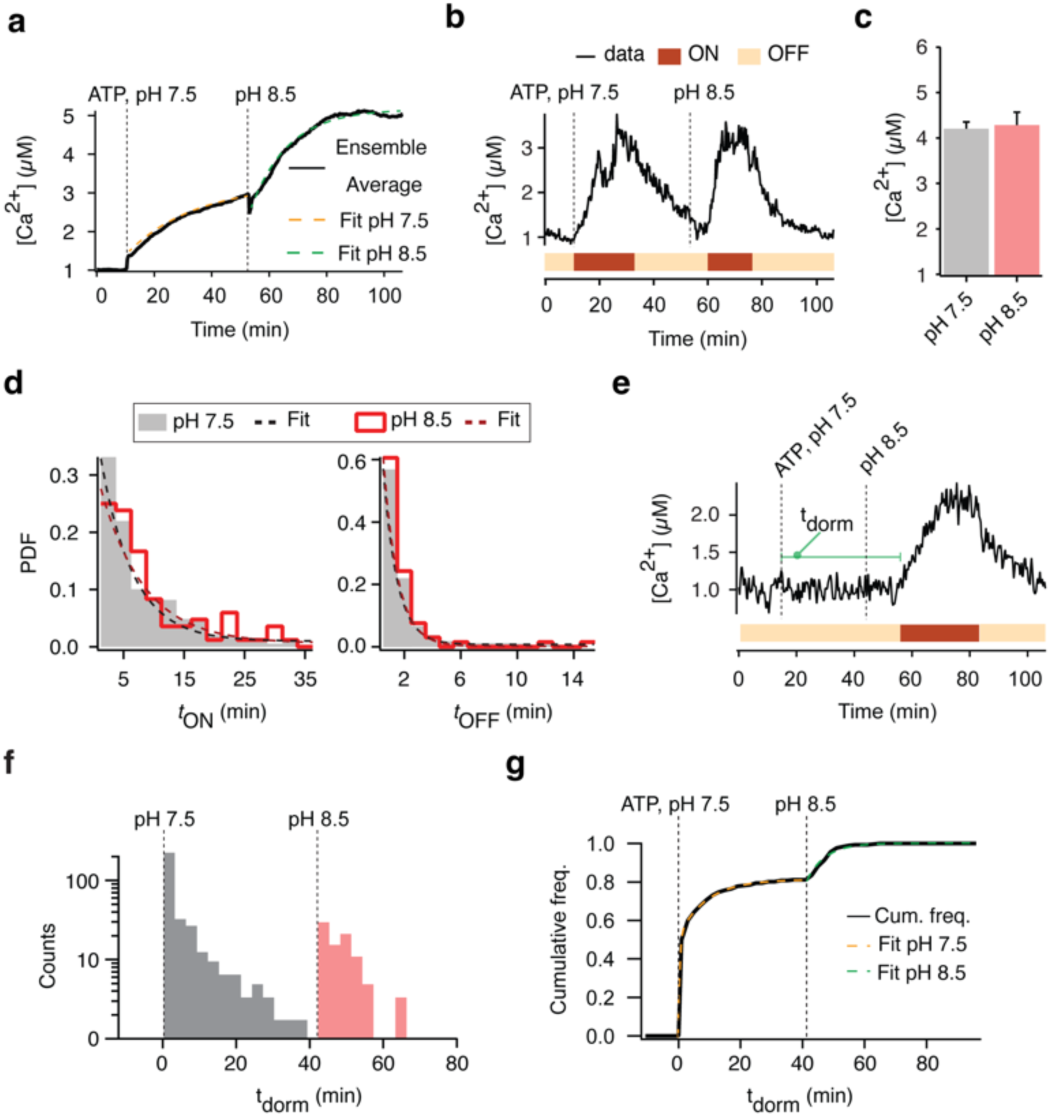
Extravesicular pH regulates the delay time until initial activation (t_dorm._), a novel mode of LMCA1, but not active and inactive modes. **a,** Ensemble average activity trace displaying Ca^2+^ upconcentration in response to ATP and an applied pH-jump from pH 7.5 to pH 8.5, and representative single vesicle traces. Data are from 178 single vesicles. *n* = 3. **b,** Single-vesicle Ca^2+^-pumping kinetics by LMCA1 that demonstrated instant activation (t_dorm_ = 0) and similar pumping activity at pH 7.5 and 8.5. c, Extraluminal pH does not affect the Ca^2+^ gradients established by single LMCA1 molecules. **d,** Histograms of active and inactive mode dwell-times for independent activity measurements of LMCA1 at pH 7.5 and pH 8.5. The histograms of t_ON_ and t_OFF_ were fitted with exponential functions to extract decay constants 1_ON_ and 1_OFF_ (1_ON, pH 8.5_ = 425 s ± 78 s, 1_ON, pH 7.5_ = 326 s ± 24 s and 1_OFF, pH 8.5_ = 79 s ± 7 s, 1_OFF, pH 7.5_ = 78 s ± 8 s). No significant pH dependence (2-sided Kolmogorov-Smirnov test) for either the active *t_on_* dwell-times (*P = 0.39*) or the inactive t_off_ dwell times (*P = 0.72*) was found. The histograms were based on 263 and 64 single-molecule traces, respectively. *n* = 3 for each pH. **e,** Single-vesicle Ca^2+^-pumping kinetics displaying delayed activation (t_dorm_). **f,** Distribution of t_dorm_ over time. The injection of ATP triggered a burst of stochastic pump activation events that plateaued after ∼30 min. The subsequent pH-jump to 8.5 induced a second burst of activation, initiating further LMCA1 activation in the remaining vesicle population displaying detectable LMCA1 activity during the experiment. **g,** Cumulative distribution of t_dorm_.

Instead, the dominant phenotype was pH-dependent recruitment of previously silent vesicles into the active switching regime. A substantial fraction of LMCA1 vesicles displayed no detectable pumping for extended periods after ATP addition at pH 7.5 yet activated robustly after the pH jump (Fig. 3e). We quantify this behavior using the delay time to first activation (*t*_dorm_), defined as the time elapsed from a triggering condition (ATP addition or the pH jump) until the first detectable Ca^2+^ pumping event. The distribution of *t*_dorm_ showed an initial burst of stochastic activation events after ATP addition that decayed and plateaued within ∼30 min, followed by a second, distinct burst of activation events immediately after shifting to pH 8.5 (Fig. 3f). The corresponding cumulative plot revealed two clear plateaus: one reached at pH 7.5 and a second reached only after alkalinization (Fig. 3g). These data indicate that extravesicular pH regulates the probability that LMCA1 exits a long-lived dormant, pre-activation off mode, thereby controlling how many molecules enter productive cycling during the experiment.

Consistent with this interpretation, once LMCA1 had already entered the on/off switching regime, pH had little detectable influence on ultraslow mode-switching kinetics. Dwell-time analysis showed that the lifetimes of the pumping and inactive modes were statistically indistinguishable at pH 7.5 and 8.5 (Fig. 3d; *P* = 0.4 and 0.7, respectively).

Together, these measurements show that pH regulation of LMCA1 primarily manifests through control of the pre-activation lifetime, modulating occupancy/escape from a long-lived dormant mode rather than tuning the intrinsic pumping rate or ultraslow switching once the transporter is active. Functionally, this implies the existence of at least two distinct “off” modes: (i) a dormant, pre-activation inactive mode that gates entry into productive cycling, and (ii) the inactive mode visited reversibly during ultraslow switching after activation.

Physiologically, it points to LMCA1 as a stress-response calcium-proton pump (Ca^2+^ out, H^+^ in), which is mostly dormant at neutral pH, but activated at alkaline cytosolic conditions in the bacterium.

### Human SERCA exhibits ultraslow mode-switching without dominant transprotein Ca^2+^ leakage

To test whether ultraslow mode-switching extends to mammalian Ca²⁺ pumps, we next examined human SERCA (hSERCA) in native ER vesicles (Fig. 4a; Fig. S16). Single-molecule measurements have traditionally relied on purified and reconstituted proteins, as performed in the present work for LMCA1. For mammalian pumps, however, purification and reconstitution can decouple the enzyme from the native organelle lipidome and proteome that shape its kinetics and regulation. For SERCA, ER microsomes represent an attractive native source. Building on our earlier single-molecule recordings of V-ATPase in intact synaptic vesicles^6^, we therefore developed a workflow to isolate biochemically intact ER-derived native vesicles from HEK293 cells (Methods; Supplementary Methods). Sequential extrusion yielded a homogeneous vesicle population with a mean diameter of ∼70 nm (Fig. S17). Vesicles were mixed with a Ca²⁺-sensitive indicator and biotinylated cholesterol, snap-frozen in liquid nitrogen, and stored at −80 °C. Immediately before experiments, samples were thawed at room temperature; the accompanying phase transition transiently permeabilized the membrane, enabling one-step loading of indicators into the vesicle lumen (Fig. 4a).

**Fig. 4.**
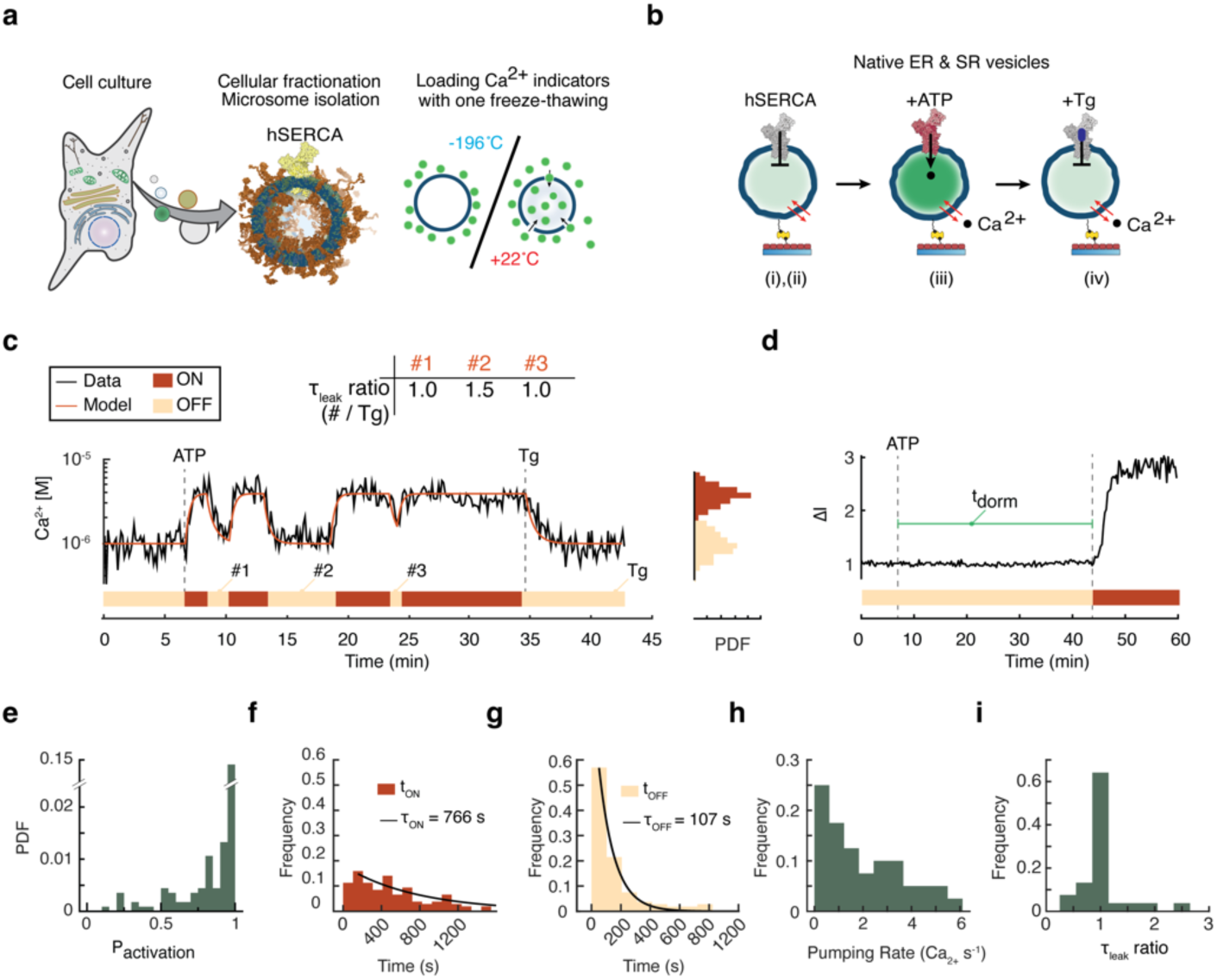
Single human endogenous SERCA pumps demonstrate delayed activation and mode-switching but do not possess a leaky mode. **a,** Illustration of the methodology developed to generate ER-derived native vesicles that contain hSERCA pumps and are loaded with calcium-sensitive fluorescent indicators in one freeze-thawing step. **b,** Vesicles containing hSERCA pumps are immobilized on a glass surface via a biotin-Neutravidin protocol (i) and permeabilized by addition of ionomycin (ii) and monitored using fluorescence microscopy. Ca^2+^ is initiated upon the addition of ATP (iii) and then is subsequently blocked upon the hSERCA-specific inhibitor Thapsigargin (Tg) (iv). Membrane permeability was tuned using ionomycin. **c,** Example kinetic trace displaying Ca^2+^ pumping activity by a single hSERCA. The pump stochastically transitions between long-lived active and inactive modes. Single-molecule kinetic traces were fit with a non-equilibrium physical model, and their leakage lifetimes during inactivity periods were monitored and compared to Ca^2+^ leakage upon addition of Tg. The leakage lifetime ratios are shown in the table and are indicative of a single leakage pathway. **d,** Example single-vesicle Ca^2+^-pumping kinetics displaying delayed activation (t_dorm_) upon addition of ATP. **e,** Histogram of activation probability shows for [Ca^2+^] = 1 μM and [ATP] = 1 mM. **f-g**, Histograms of active and inactive mode dwell times are fitted with single exponential functions. **h,** Single hSERCA pumping rates as calculated by the physical model. **i,** Histogram of leakage lifetime ratios. The lifetime of each stochastic event is normalized to the lifetime of the last event triggered by Tg inhibition, on a per-vesicle basis. The prominent peak at 1_leak_ = 1 suggests the existence of a single leakage pathway, passive leakage through the membrane.

To isolate and positively identify hSERCA activity at the single-vesicle level, we used a four-step protocol (Fig. 4b): (i) immobilization on a NeutrAvidin functionalized surface enabled extensive washing to remove non-encapsulated fluorophores and time-lapse fluorescence microscopy imaging; (ii) controlled ionomycin permeabilization of the otherwise Ca^2+^-intact vesicles (Fig. S18) established the self-regenerating measurement regime; iii) ATP addition revealed the subset of vesicles containing an active Ca²⁺ pump; and (iv) addition of thapsigargin^42^ (Tg) allowed us to attribute the activity specifically to SERCA, distinguishing it from e.g. plasma-membrane Ca²⁺-ATPases (Fig. 4b, c; Fig. S19a).

Similar to LMCA1, single-vesicle traces revealed delayed activation and stochastic switching between long-lived pumping and inactive modes (Fig. 4c, d, and Fig. S19b-j). Across the population, the probability of occupying the pumping mode varied broadly from vesicle to vesicle (Fig. 4e), indicating stable molecule-to-molecule heterogeneity in the balance between pumping and inactivity on the timescale of our recordings. For molecules exhibiting mode-switching, the characteristic lifetimes were τ_ON_ = 766 ± 90 s and τ_OFF_ = 107 ± 15 s (Fig. 4f, g), resulting in an average *P*_ON_ ∼ 0.9, very similar to LMCA1 (*P*_ON_ ∼ 0.8). Fitting each trace with the non-equilibrium physical model allowed us to deconvolve pumping from inactive modes, leakage, and indicator response (Fig. 4c, h, Supplementary information, Table S1) and thus calculate the average single-molecule Ca^2+^ pumping rates to be *I*_SERCA_ = 2.4 ± 0.3 Ca^2+^ s^-1^, in excellent agreement with previous studies^43^. Together, these data demonstrate that ultraslow mode-switching is a bona fide single-molecule phenotype of hSERCA in native membranes.

In contrast to LMCA1, we did not detect evidence for a prominent transprotein Ca^2+^ leakage mode in hSERCA in native lipid ER microsomes. Specifically, the kinetics of Ca^2+^ efflux during inactive periods closely matched those observed after Tg inhibition in the majority of vesicles, consistent with a single dominant leakage pathway (Fig. 4c, i). Thus, while both bacterial and human Ca^2+^-ATPases exhibit ultraslow switching between pumping and inactivity, their access to uncoupled leaky states differs markedly.

### ATP and Ca^2+^ tune hSERCA activation probability by modulating dormant-mode occupancy

Because delayed activation was the sole regulatory pathway for LMCA1 (both mode lifetimes and pumping rates exhibited remarkable pH stability), we next asked whether the dormant mode also serves as a regulatory checkpoint for mammalian hSERCA. Since cytosolic Ca²⁺ is the dominant physiological regulator of hSERCA activity, and nucleotide availability further determines pump flux, we performed experiments in which ATP and Ca^2+^ concentrations were varied by sequential injections while monitoring the emergence of pumping activity in individual vesicles (Fig. 5a).

**Fig. 5.**
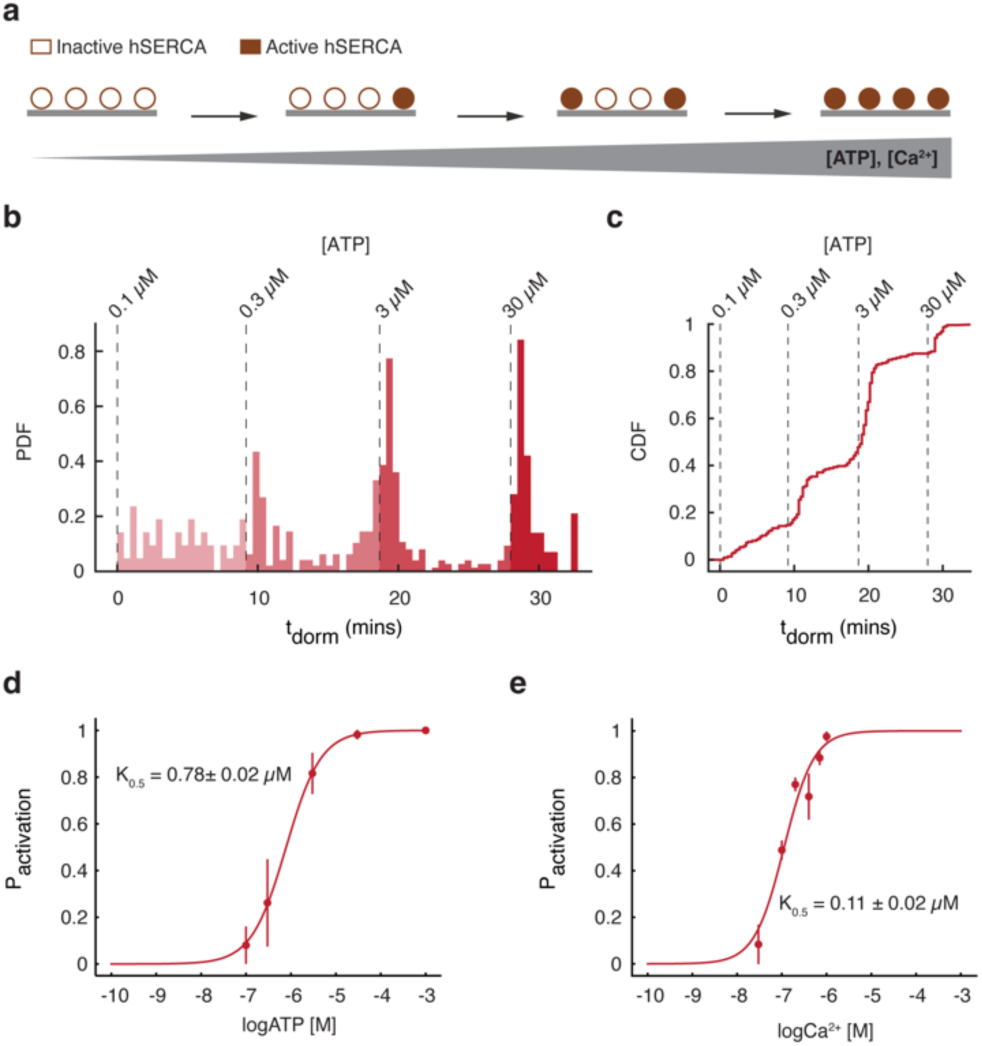
Ca^2+^ and ATP regulate the dormant-mode occupancy and thus the activation probability of hSERCA. **a,** Illustration of stochastic activation of hSERCA in single vesicles with increasing concentrations of either ATP or Ca^2+^. **b,** Distribution of delayed activation lifetimes (t_dorm_) over time and over a range of ATP concentrations showcases bursts of activating vesicles upon addition of each ATP injection. **c,** Cumulative distribution of t_dorm_. **d, e,** Titration curves of hSERCA activation probability over a range of ATP (d) and Ca^2+^ (e) concentration show that the fraction of active hSERCA molecules increases with increasing catalytic or ionic substrate concentration.

Quantifying the pre-activation dormant delay time (*t*_dorm_) revealed bursts of newly activating vesicles after each ATP injection, with the rate of new activations decaying over time (Fig. 5b). hSERCA molecules that activated early on retain the same activity level during the half-hour-long experiments (Fig. 4c, S19), demonstrating that there is no significant depletion or autohydrolysis of ATP during our measurements (see also Fig. S20). Cumulative distributions of *t*_dorm_ across independent experiments showed that both ATP and Ca^2+^ systematically shift the probability of activation (Fig. 5c). Consistently, titration curves demonstrated that the fraction of vesicles that become active increases with ATP and with Ca^2+^ concentration (Fig. 5d, e). These observations reveal a novel regulatory axis through which ATP and Ca^2+^ can regulate hSERCA activity by controlling the occupancy and escape kinetics of the dormant state, thereby tuning the probability of entering productive pumping.

### Discussion, conclusions

Long-duration single-molecule flux measurements established mode-based regulation in two proton pumps^5,6^, but the extent to which this behavior generalizes beyond proton transport has remained unresolved. Here we introduce two complementary methodological advances: (i) a self-regenerating single-vesicle flux assay that renders zeptoampere Ca^2+^ currents continuously observable by tuning Ca^2+^ permeability with ionophores, and (ii) a workflow enabling single-molecule measurements of endogenous pumps in native lipid vesicles. Using these methods, we show that Ca^2+^-ATPases exhibit ultraslow stochastic switching between functional modes that persist for minutes. This finding extends the mode-switching paradigm to the alternating-access mechanisms characteristic of Ca^2+^ transport^44^, supporting an emerging view that transporter regulation can be implemented by controlling *access* to productive cycling^5,6^.

Beyond suggesting mode generality across ion classes, our observations recast how Ca^2+^ pump regulation can be conceptualized at the molecular level. Once a pump is in the pumping mode, the observed activity is compatible with progression through the classical Post-Albers cycle^26^ (*active* mode); however, it can stochastically exit the cycle and enter extended periods of inactivity^5^ (*inactive* mode). Crucially, we identify a pre-activation *dormant* mode that gates entry into productive cycling and governs regulation by pH, ATP, and Ca^2+^. Functionally, the activation from dormancy complements other autoinhibitory mechanisms based on, e.g., calcium-calmodulin-, lipid-, or alpha-synuclein-sensitive autoinhibition^45,46^. The presence of an uncoupled Ca^2+^ leak (in LMCA1 but not in hSERCA under our conditions) raises the possibility that *leaky* modes may contribute to physiological Ca^2+^ leak, heat generation^47^, or stress responses^48^, or represent rare malfunction states^49^, which will be important to address in future work.

Ca²⁺ differs from most transported solutes in that it is both an essential cofactor and a ubiquitous second messenger whose concentrations are controlled by feedback, thresholds, and spatial microdomains^28^. More broadly, our findings suggest a regulatory logic for Ca²⁺ handling that is naturally suited to non-linear cellular physiology. Because Ca²⁺ homeostasis depends on rapid transitions between release and sequestration and is often organized into local compartments, probabilistic control of pump availability via dormant and inactive mode occupancy provides a mechanism to tune Ca²⁺ dynamics by altering activation latency and duty cycle. In this regime, regulatory inputs can reshape when pumping begins and how consistently it proceeds across individual pumps, without measurably altering Post-Albers cycle kinetics once pumping is underway. Such probability-gated control explains why substantial functional variability may coexist with apparently stable ensemble-average transport rates.

We anticipate that the methods presented here could, in principle, be extended to monitor the transport of other membrane-impermeable ions (e.g., Na^+^ or K^+^) by human endogenous or exogenous pumps and secondary ion-coupled transporters, helping us understand how leak pathways and *ultraslow* mode-switching shape function over physiological time scales.

## Methods

### Chemicals

L-α-Phosphatidylcholine (soybean, Type II-S, 14-23% choline basis, (Sigma-Aldrich, Cat. No. P5638). 18:1 Biotinyl-Cap-PE (Avanti Polar Lipids, Cat. No. 870273). 1,2-dioleoyl-sn-glycero-3-phosphoethanolamine headgroup labeled with ATTO655 (ATTO655-DOPE, ATTO-TEC GmbH, Cat. No. AD 655-161). Cal-520 potassium salt (AAT Bioquest, 21140). Fluo-5N, pentapotassium salt, cell impermeant (ThermoFisher Scientific, Cat. No. F14203). Detergent n-Dodecyl-β-D-maltoside (DDM, Glycon Biochemicals, Cat. No. D9702). Unless indicated otherwise, all other chemicals and reagents were obtained from Sigma-Aldrich (Brøndby, Denmark).

### Liposome preparation for LMCA1 reconstitution

Unilamellar liposomes were prepared by manual extrusion. Briefly, chloroform stocks of phospholipids (12.7 μmol, L-α-phosphatidylcholine), DOPE-biotin (38 nmol; 0.3 mol%), and ATTO655-DOPE (12.7 nmol; 0.1 mol%) were combined into a round-bottom flask and dried under vacuum for ≥1 h. The resulting lipid film was rehydrated in 667 µL of reconstitution buffer (20 mM Tris-HCL, pH 8.5, 200 mM KCl, 3 µM EGTA) supplemented with 1 mM Cal-520, dissolved by vortexing in the presence of a glass pearl, and passed 11 times through two 0.2 μm size nucleopore polycarbonate membranes mounted in a mini-extruder (Avanti Polar Lipids). Vesicles were subjected to five freeze-thawing cycles (liquid nitrogen/ 50°C water bath) followed by a second extrusion through two 0.2 µm membranes as stated above. The resulting vesicles were kept at 4°C and used within 1 week.LMCA1 protein expression, purification and bulk ATPase activity A pET-22b:LMCA1-10xHis construct^50^ was transformed into chemically competent *E. coli* C43 (DE3) cells and plated on LB agar plates containing 100 µg/ml ampicillin. Overnight starter cultures (20 mL LB, 100 µg/mL ampicillin) were inoculated from 3-5 colonies and usedto seed 2 liters of LB supplemented with ampicillin. Culture wered grown at 37°C until the optical density at 600 nm (OD600) reached 0.6-0.8. Expression was induced with 1 mM isopropyl β-D-1-thiogalactopyranoside, and the temperature was decreased to 20°C. Cells were grown for 20 h and harvested by centrifugation (8000 g, 10 min). The resulting pellets were weighed and resuspended to 20% (w/v) in buffer A (50 mM Tris-HCl pH 7.6, 200 mM KCl, 20% v/v glycerol). Cells were lysed by high-pressure homogenization (15,000 psi, four passes) in buffer A supplemented with 5 mM β-mercaptoethanol, 1 mM phenylmethylsulfonyl fluoride (PMSF), 1 Complete protease-inhibitor tablet (Roche) per 8 L suspension, and 1 μg/ml DNase I. Cell debris and aggregates were removed by centrifugation (27,000 × *g*, 45 min). Membranes were isolated by centrifugation (235,000 × *g*, for 1.5 h), washed in a high-salt buffer B (20 mM Tris-HCl pH 7.6, 1 M NaCl, 20% v/v glycerol, 1 mM MgCl_2_, 5 mM β-mercaptoethanol), centrifuged again (235,000 × *g*, 1.5 h), and subsequently solubilized in a low-salt buffer B (200 mM KCl) with 1% w/v DDM at 4°C for 1 h. Aggregates were removed by centrifugation (235,000 × *g*, 1.5 h). Solubilized protein was purified by Ni²⁺-affinity chromatography using a 5 ml pre-packed HisTrap HP column (GE Healthcare) equilibrated in buffer C (20 mM Tris-HCl pH 7.6, 200 mM KCl, 20% v/v glycerol, 1 mM MgCl_2_, 5 mM β-mercaptoethanol, 0.25 mg/ml C_12_E_8_, 50 mM imidazole). After washing with five column volumes, LMCA1 was eluted with 150 mM imidazole. Protein purity and Ca²⁺-dependent ATPase activity were previously validated ^50^.

### LMCA1 reconstitution in proteoliposomes and characterization

Purified LMCA1 was reconstituted into preformed liposomes as described previously^5^. Briefly, 50 µg of purified protein was mixed with 220 µL reconstitution buffer containing 1 mM Cal-520, 200 µM EGTA, 52 mM octylglucoside and liposomes (10 g L^-1^). The mixture was vortexed and incubated for 30 min at room temperature with 100 mg of wet Bio-Beads (Bio-Rad Laboratories) under overhead rotation to remove the detergent. Protein-free vesicles were prepared accordingly by replacing purified protein with reconstitution buffer. Vesicles were then mixed in a 1:1 ratio (v/v) with a concentrated dye solution (20 mM Cal-520, 20 mM Tris-HCL, 200 mM KCl, 53 µM EGTA, pH 8.5) and flash-frozen in liquid nitrogen. The vesicle sizes were calculated as described previously^51^. Briefly, the number of ATTO655-DOPE fluorophores incorporated into the membrane is proportional to the liposomal surface area (*A_liposome_*) and is related to the radius (*R_liposome_*) through equation (1), thereby allowing for the conversion of fluorescence intensities into physical sizes.

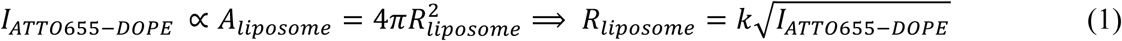

Where (*k*) is a calibration factor, which we determined as follows: The vesicles were characterized using cryogenic transmission electron microscopy (Cryo-TEM), and their diameters were measured using custom software developed in MATLAB. The mean diameter was determined by applying a gaussian fit to the resulting size histogram (Fig. S1). The mean fluorescence intensity of the diffraction-limited vesicles, as determined by fluorescence microscopy, was then correlated with the mean radius, as determined by Cryo-TEM, to calculate *k*. After the determination of *k*, all fluorescence intensities were converted to radii using equation (1).

### Cell Culture

Human embryonal kidney cells (HEK293) were cultured in DMEM media supplemented with 10% fetal bovine serum (Gibco, Thermo Fisher Scientific), 100 units/ml Penicillin and 100 µg/mL streptomycin (Gibco, Thermo Fisher Scientific) at 37°C in a humidified atmosphere with 5% CO_2_.

### Endogenous ER vesicle isolation

Cells were cultured as described above for three days, after which cell media was gently rinsed out three times with Base buffer (20 mM Tris-HCl, 200 mM KCl, 2 µM EGTA, pH 7.4). Cells were harvested in Base buffer using a cell scraper and spun down at 300 x g for 10 min. The cell pellet was re-suspended in fresh Base and cells were lysed by aspiring and ejecting the cell suspension through a 27G Fine-Ject needle (27G, L x 1/2’’, Henke Sass, Wolf) 20 times. The lysate was spun again at 300 x g for 10 min to pellet out any residual unlysed cells and the supernatant was subjected to subcellular fractionation starting with centrifugation at 800 x g for 10 min, followed by centrifugation at 10,000 x g for 10 min and ending with ultracentrifugation for 60 min at 100,000 x g_avg_ (TLA-100.3, k-factor 62, Beckman Coulter). SThe resulting microsome-enriched pellet was resuspended in fresh Base buffer and subjected to serial extrusion using a mini-extruder (Avanti Polar Lipids) through polycarbonate filter membranes with pore sizes of 5, 1, 0.8, 0.4, 0.2, and 0.1 µm (Nucleopore Track-Etched Membranes, Whatman), in this order, with 19 times passes per membrane. The entire procedure was performed on ice, using thoroughly chilled buffers. See also Fig. S16.

### SDS-PAGE and Western Blotting

Whole cell lysates and microsome lysates were generated by the addition of RIPA buffer (Cell Signaling Technologies) followed by sonication, 3 x 5 minutes in ice bath with vortexing in between rounds of sonication, ending with a centrifugation for 15 minutes at 14 000 g. Protein concentration was determined using a BCA protein assay kit (Thermo Fisher Scientific) as per manufacturer’s instructions. For SDS-PAGE, 20 µg protein from each lysate were loaded onto 4-12 % Bis-Tris gels under reducing conditions. Gels were blotted onto PVDF membranes using the Trans-Blot Turbo transfer system (Bio-Rad). Membranes were blocked with EveryBlot blocking buffer (Bio-Rad) for 10 min and then probed with primary antibodies against SERCA (Recombinant Anti-SERCA2 ATPase antibody [EPR9393], ab137020, Abcam) diluted 1:1,000 in blocking buffer with gentle rocking over night at 4°C. Membranes were washed with TBS buffer containing 0.1% Tween-20 and then incubated with HRP-conjugated secondary antibodies (Goat Anti-Mouse IgG H&L (HRP) preadsorbed, ab97040, Abcam) diluted 1:10,000 for 90 min in blocking buffer at room temperature with gentle rocking. Blots were developed using SuperSignal West Pico PLUS Chemiluminescent Substrate (Thermo Fisher Scientific) and imaged. See also Fig. S17.

### Sensor loading into ER vesicles

Extruded vesicles were loaded with Cal-520 (Cal-520 potassium salt, AAT Bioquest) by combining 15 µl of vesicle sample (∼30 µg worth of total protein) with 5 mM of Cal-520 dye and 50 µM of EGTA in Base at a final volume of 20 µL. Dye loading into the vesicular lumen was achieved through freeze-thawing, whereby the sample/dye mixture is submerged into liquid nitrogen until frozen and then either stored at −80°C for later use, or thawed on ice directly for immediate downstream applications.

### Surface preparation and immobilization of vesicles

Surfaces were prepared according to previously published protocols^29–31,51^. Briefly, flow cells were prepared by assembling sticky slides VI 0.4 (Ibidi) and glass slides (thickness 170 ± 10 μm). Prior to assembly, the glass slides passed sequential sonication cycles in 2% (v/v) Helmanex III and Milli-Q water and subsequently rinsed in 70% (v/v) ethanol and 99% (v/v) methanol. The glass slides were dried under nitrogen flow, plasma etched for 3 minutes to remove impurities and then immediately assembled with the sticky slide. The surface was functionalized by incubation with mixture of 1.0 g/L BSA-biotin/BSA solution (1:10) for 10 min. Unbound BSA-biotin/BSA was removed by flushing the flow chambers with 15mM HEPES buffer. Subsequently, the glass surface was incubated with 0.1 g/l Neutravidin (Life Technologies) for 10 min. Unbound Neutravidin was removed by flushing with immobilization buffer (20 mM Tris-HCl, 200 mM KCl, 60 nM valinomycin, 3 μM EGTA). The immobilization buffer always had matching ionomycin concentration to the experiment for which the vesicles were subsequently used. A vesicle solution was prepared by diluting 0.2 μL of the sample (7.5 g L^-1^) in immobilization buffer to a final concentration of 7.5 mg L^-1^. The vesicle solution was flushed into the flow chamber and incubated until the density of immobilized vesicles reached approximately ∼1000 in an 81.92 µm × 81.92 μm field-of-view (FOV). Unbound vesicles and free-floating dye was removed by flushing repeatedly with transport buffer (20 mM Tris-HCl, 200 mM KCl, 1mM MgCl_2_, 60 nM valinomycin, 3 μM EGTA, 0 or 1 μM CaCl_2_, pH 7.5).

For native vesicles, an additional incubation for 10 minutes with 0.1 mg/ml Cholesterol-PEG-Biotin (PG2-BNCS-2k, Nanocs) was implemented, followed by a wash step to eliminate unbound compound. A vesicle solution was prepared by diluting either 0.2 μL of synthetic vesicles (7.5 g L^-1^) in immobilization buffer to a final concentration of 7.5 mg L^-1^, or by diluting native vesicles at a 1:25 ratio in immobilization buffer. The vesicle solution was flushed into the flow chamber and incubated until the density of immobilized vesicles reached approximately ∼1000 in a 81.92 µm × 81.92 μm field-of-view (FOV). Any inconsistencies in native vesicle concentrations were thus accounted for by incubation duration. Unbound vesicles and free-floating dye was removed by flushing repeatedly with transport buffer.

### Image acquisition

§All fluorescence microscopy was performed on a commercial Olympus total internal reflection fluorescence microscope (Olympus Europa). We used the EMCCD camera iXon 897 (Andor Technology). Excitation of fluorophores was achieved using Olympus Cell solid-state lasers with excitation wavelengths of 488 nm (Ca^2+^ indicator, Cal-520) and 640 nm (membrane dye, ATTO655-DOPE). The membrane dye was imaged in wide-field mode to ensure illumination of all membrane-incorporated fluorophores, while the Ca^2+^ indicator channel was imaged in TIRF-mode to ensure high SNR of kinetic recordings with minimal background fluorescence. The microscope was equipped with an Olympus total internal reflection fluorescence UApoN ×100, 1.49 NA, oil immersion objective. The microscope was able to maintain focus for extended periods of time using the continuous function of the Zero Drift Correction module built into the microscope. An emission filter with bandpasses 562.5/62.5 nm and 712.5/47.5 nm and beamsplitter bandpasess 563.5/69.5 nm and 708.5/61.5 nm were used to block fluorophore excitation. Image acquisition software linked to the microscope and used throughout this article was for the most part Olympus xcellence rt (version 2.0) and to a lesser extent Olympus cellSens (version 3.2). Replicates of measurements for each condition were taken on the same sample measured repeatedly throughout the present manuscript. Images were acquired in the format of 512 × 512 pixels, each pixel corresponding to 160 nm sample length and a bit depth of 16.

### Cryo-Electron microscopy

EM grids (Lacey Carbon grids, Quantifoil) were glow-discharged (Leica EM MED020, Leica), mounted in a Vitrobot (Vitrobot Mark IV, FEI), and a sample volume of 3 µl was deposited unto them followed by automated blotting and vitrification by plunging the grid into liquid ethane. Images were acquired using a transmission electron microscope (Tecnai G2 20 TWIN, FEI) and exported as tif-files which were then manually analyzed using ImageJ (version 1.52p) or custom, in-house, software developed in MATLAB.

## Acknowledgements

We thank M. Grabe for providing the MatLab code for fitting the non-equilibrium physical model. This work was supported by the Lundbeck Foundation (Professorship grant R441-2023-360) and the Novo Nordisk Foundation (NNF17OC0028176).

## Author contributions

D.S. conceived the strategy and was responsible for project management. D.S. and E.K. supervised the project. M.P.M. and A.C. designed research. M.P.M. developed the LMCA1 assay and performed all experiments with the help of J.H. A.C. developed the SERCA assay and performed all experiments with the help of A.N. and M.I.. C.G.S. developed kinetic modelling and was the principal software developer. M.P.M., A.C. and A.N. performed data analysis with the help of M.I. M.D. purified LMCA1 under the supervision of M.K. and P.N.. B.H.J and I.L.J reconstituted LMCA1 under the supervision of T.G.P.. P.A.P. assisted with ER purification. D.S. and E.K. wrote the main text. M.P.M., A.C., C.G.S. and E.K. wrote supplementary methods, materials and information. All authors discussed the results and commented on the manuscript.

## Supplementary methods

### Measurement of LMCA1 Ca^2+^ pumping kinetics in low-permeability vesicles

The fluorescence intensity of the Ca^2+^ indicator of surface-tethered vesicles was initially monitored continuously for 5 min (7.5 frames min^-1^) to establish a baseline in the absence of ATP (20 mM Tris-HCl, 200 mM KCl, 1mM MgCl_2_, 60 nM valinomycin, 3 μM EGTA, 200 nM ionomycin, 1µM CaCl_2_, pH 7.5). LMCA1 activity was initiated by the injection of 1 mM ATP (20 mM Tris-HCl, 200 mM KCl, 1mM MgCl_2_, 60 nM valinomycin, 3 μM EGTA, 200 nM ionomycin, 1µM CaCl_2_, 1mM ATP, pH 7.5) and the fluorescence intensity of the Ca^2+^ indicator was monitored continuously for 35 min (7.5 frames min^-1^). To detect ultra-slow Ca^2+^ pumping in some experiments, we subsequently flushed ATP out of the system, to immediately stop Ca^2+^ pumping activity and monitored for an additional 40 min (1 frame min^-1^) to confirm that Ca^2+^ did not leak out of the vesicles over the timescale of an experiment (Fig. 1c, Fig. S6).

### Calcium permeability assay

Surface-immobilized vesicles were pre-incubated with varying [ionomycin] for at least 15 min (20 mM Tris-HCl, 200 mM KCl, 1mM MgCl2, 60 nM valinomycin, 3 μM EGTA, 0-2,000 nM ionomycin, pH 7.5) and subsequently exposed to a 2 mM Ca^2+^ gradient (20 mM Tris-HCl, 200 mM KCl, 1mM MgCl_2_, 60 nM valinomycin, 3 μM EGTA, 2 mM CaCl2, 0-2,000 nM ionomycin, pH 7.5). We adjusted the experimental settings to match the timescale of the equilibration kinetics. Framerates of 1.00, 1.89, 13.0, and 17.5 s^-1^ were used for 2, 20, 200 and 2,000 nM ionomycin, respectively. Single-vesicle Ca^2+^ equilibration kinetics were monitored as intensity increases as Ca^2+^ leaked into the vesicle and eventually saturated the sensor signal (ΔI = 1, Supplementary methods). Each trace was fitted to a sigmoidal function of the form:

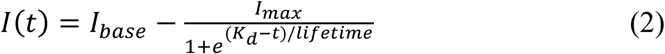

To extract lifetimes as a function of ionomycin concentration.

### Transport assay for measuring long-lived mode-switching by single LMCA1

LMCA1 activity was initiated by the injection of 1 mM ATP (20 mM Tris-HCl, 200 mM KCl, 1mM MgCl2, 60 nM valinomycin, 3 μM EGTA, 200 nM ionomycin, 1µM CaCl2, 1mM ATP, pH 7.5) to the surface-tethered vesicles, and the fluorescence intensity of the Ca^2+^ indicator was monitored continuously for either 35 min (7.5 frames min^-1^) or 240 min (1 frame min^-1^). When studying the transprotein leaky state of LMCA1, we blocked the pumping activity by injecting 0.1 mM vanadate after 35 min and then monitored the passive Ca^2+^ efflux for 25.2 min (7.5 frames min^-1^). When studying the effect of pH on LMCA1 Ca^2+^ pumping kinetics we recorded for a total of 106.2 min (3.75 frames min^-1^), and introduced a pH-jump from pH 7.5 to pH 8.5 (20 mM Tris-HCl, 200 mM KCl, 1mM MgCl_2_, 60 nM valinomycin, 3 μM EGTA, 200 nM ionomycin, 1µM CaCl2, 1mM ATP, pH 8.5) after 53.1 min and subsequently monitored for an additional 53.1 min.

### Transport Assay for measuring long-lived mode-switching by single hSERCA

Following vesicle immobilization and sample mounting on a TIRF microscope, Assay buffer (Base buffer supplemented with 60 nM Valinomycin, 1 µM Ionomycin, 1 mM MgCl2, and 1 µM CaCl2) was introduced into the flow chamber and allowed to equilibrate for five min to establish baseline conditions before commencing image aquisition. Transporter activity was initiated by the replacement of MgCl2 in the Assay buffer with 1mM MgATP (Mg^2+^/ATP Activating Solution, BML-EW9805-0100, Enzo Life Sciences). Pump inhibition was facilitated by the addition of Assay buffer containing ATP, supplemented with 60 nM Thapsigargin (1138, Bio-techne). The transport assay spanned a duration of 42 min and 40 s. The first 6 min and 40 s were dedicated to baseline recording, then 28 min of activity recording, and lastly 8 min of inhibition recording. The experiment ended with a separate photobleaching recording under saturating conditions which were achieved by flushing the chamber with Saturating buffer (Base buffer supplemented supplemented with 2 µM ionomycin and 2 mM CaCl_2_). Baseline, transport activity, and inhibition were captured in one continuous recording. Photobleaching was recorded after 5 min equilibration with Saturating buffer with identical microscope settings as those used for the transport

### Additional image acquisition for corrections and calibrations

At the end of each kinetic measurement, a saturation buffer (20 mM Tris-HCl, 200 mM KCl, 1mM MgCl2, 60 nM valinomycin, 3 μM EGTA, 2 mM CaCl2, 2 μM ionomycin, pH 7.5 or pH 8.5) was flushed into the flow chamber to collapse any established Ca^2+^ gradients and saturate the response of the encapsulated Ca^2+^ indicator. The system was allowed to equilibrate for 10 min before a photobleaching movie was recorded in the same FOV. The photobleaching traces were used for local correction^52^ of single-vesicle activity traces, Fig. S5 and Supplementary methods, and to determine the dynamic range of each vesicle, which was subsequently used for semi-local calibration of activity traces from intensity to [Ca^2+^] (Fig. S3 and Supplementary methods). The vesicles were imaged in the membrane channel before- and after all buffer exchanges to ensure that we could identify and exclude vesicles attaching or detaching from the surface during the experiment (Supplementary methods). Finally, the vesicles were removed from the surface by flushing with a solution of 0.2% Triton X-100 (w/v), and local illumination profiles were acquired in both channels in the presence of soluble fluorophores (Cal-520 for the Ca^2+^ indicator channel or Cy5 for the membrane channel). The respective profiles were used to correct all fluorescence microscopy data for uneven illumination of the FOV.

### Image analysis

Data from fluorescence microscopy experiments were analyzed using customized software developed in MATLAB. All acquired images were initially corrected for dark camera noise by subtracting an averaged image sequence acquired with the camera shutter closed. Image sequences were subsequently corrected for variations in the illumination profile, by dividing through by normalized, locally acquired control images of the illumination profiles of the membrane channel and Ca^2+^ sensor channels, respectively. A drift correction algorithm was implemented to compensate for xy-drift during acquired time sequences and to align all image sequences between the membrane- and sensor channel^53^. Areas that were out-of-focus or contained artefacts were cropped out and excluded from further analysis. A single particle localization algorithm^54^ identified the vesicle positions within the FOV based on peaks in the intensity profile. Particles spatially separated by less than 5 pixels were excluded from the analysis to avoid significant intensity contributions from neighboring particles. Particle positions that colocalized between the membrane- and sensor channel within no more than 2 pixels were used for further analysis. Fluorescence intensities recorded in the sensor channel were extracted by applying a 5×5 pixel region-of-interest (ROI) around the identified particle centers and summing up the intensity values of the included pixels. Single-particle kinetics were extracted by ROI summation of every image throughout the recorded time series. The membrane intensities were quantified by 2D Gaussian fitting of the intensity profiles and extracting the integrated fluorescence intensity.

We defined Δ*I* = (*I* – *I*_base_)/(*I*_max_ - *I*_base_) as the relative change in fluorescence response between the baseline intensity (*I*_base_) and the intensity at saturating [Ca^2+^] (*I*_max_), which is the standard representation of fluorescence microscopy intensity data in this work. In this representation, the data points take a value between 0 and 1.

### Post-processing and data filtering

We initially applied filters to the acquired microscopy data based on the sensor- and membrane intensity of individual vesicles: Aggregates were excluded by threshold filtering particles with high membrane signals. Vesicles that detached from- or attached to the surface during the experiment were identified and excluded based on membrane intensities ratios recorded before and after kinetic measurements. Vesicles with too low amounts of encapsulated Ca^2+^ indicator were excluded based on *I*_max_, determined in the presence of saturating Ca^2+^ concentrations (See also Supplementary methods and Fig. S2). Activity traces that passed these initial filtering steps were corrected for photobleaching, using their corresponding photobleaching trace (See also Fig. S5). Briefly, photobleaching traces were fitted with exponential functions and then normalized to the initial intensity. The entire image sequence for each vesicle was then divided through by its corresponding photobleaching fit. We found that the photobleaching rates varied from vesicle to vesicle, stressing the importance of implementing local photobleaching corrections. The source of this heterogeneity is likely dominated by variations in the illumination profile across the FOV, thus exposing vesicles to varying laser intensities. A minor fraction of vesicles (>1%) displayed characteristic self-quenching due to “overloading” with the Ca^2+^ indicator. Such particles were identified manually based on their photobleaching trace and excluded from further analysis. After correcting for photobleaching, the activity traces were threshold filtered based on the dynamic range of their responses to changes in luminal [Ca^2+^] (*I_max_/I_base_*). Particles with relative responses < 2 were excluded from further analysis. Vesicles with active transporters were identified by threshold filtering the signal-to-noise (SNR) of the kinetic traces. The SNR was determined using the maximum value of standardized intensity (*I* – mean(*I*_base_)/std(*I*_base_)) defined for each trace as the maximum normalized intensity reduced by one and divided by the normalized standard deviation of the baseline (*SNR = (I_norm,max_ - 1)/σ_norm,base_*).

### Calibration from intensity to Ca^2+^ concentration inside vesicles

Single-molecule activity traces were converted into absolute [Ca^2+^] using a semi-local calibration method (See also Fig. S3). We recorded single vesicle dose-response curves by imaging Ca^2+^-equilibrated vesicles in the presence of 12 different calibration buffers (20 mM Tris-HCl, 200 mM KCl, 1mM MgCl_2_, 3 μM EGTA, 60 nM valinomycin, 2 μΜ ionomycin, pH 7.5) containing varying [Ca^2+^] (from 0.05 to 2000 μM CaCl_2_). Ionomycin and valinomycin facilitated a free influx of Ca^2+^ and prevented the buildup of cross-membrane electrical gradients. We acquired and averaged a stack of 3 images for every [Ca^2+^] to increase the SNR and calculate the s.d. for every measurement point. Dose-response curves were recorded as *n*=3 independent replicates. We fitted single vesicle dose-response curves to sigmoidal functions of the form:

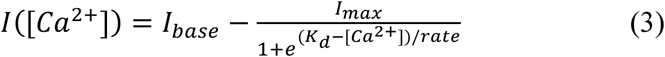

Where *I* is the raw intensity, *I*_base_ is the lower asymptote, I_max_ is the upper asymptote, *K_d_* is the dissociation constant of the Ca^2+^ indicator, and rate is describing the steepness of the sigmoidal curve. 90 ± 1.0% of the single vesicle calibration curves could be fitted with an adjusted *r*^2^ > 0.97 (*n*=475 single vesicles). Single vesicle analysis revealed a narrow distribution of *K_d_* (4.44 ± 1.6 μM) and rate (0.316 ± 0.0316 M), in contrast to the dynamic range (*I*_max_/*I*_base_) which spanned from ∼1 to > 40-fold (Fig. S3). For this reason, we combined the globally determined values for Kd and rate with locally acquired Imax for the conversion from intensity to luminal [Ca^2+^]. We normalized all activity traces to units of ΔI, thus defining the intensity range for each vesicle to go from 0 (I_base_) to 1 (I_max_). Therefore, equation (3) could be simplified to:

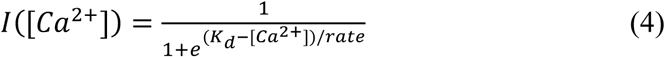

for semi-local calibrations. Solving equation (4) for [Ca^2+^], we converted all intensity data points to luminal [Ca^2+^].

### Chelation of Ca^2+^ contaminants

Contaminants of free Ca^2+^ ions were chelated by adding an equivalent concentration of EGTA to all buffers. The amount of EGTA required to chelate all Ca^2+^ ions was determined using a Horiba Jobin Yvon FluoroMax-4 spectrofluorometer. The temperature was kept at 25°C for all measurements, using a Wavelength Electronics LFI-3751 thermoelectric temperature controller. Ca^2+^ indicator, Fluo-5N, was diluted to a final concentration of 2.6 µM in buffer (20 mM Tris-HCl, 200 mM KCl, 1mM MgCl2, pH 7.5) and added to a 1500 μL quartz cuvette. The solution was continuously stirred with a magnetic micro-stirrer while the EGTA concentration was increased stepwise. The solution was excited with an excitation wavelength of 491 nm with a slit width of 2 nm. Emission was recorded at 516 nm a slit width 2 nm and an integration time of 0.1 seconds. Between each independent experiment (*n*=5), the cuvette and magnet were rinsed sequentially 5 times with acetone, MilliQ-water and methanol, and dried under nitrogen flow. See also Fig. S4.

### Quantification of lumen-dye concentration

Concentrations of encapsulated fluorescent indicator ([Cal-520], “lumen-dye”) were obtained by assuming that the luminal concentration could not exceed the concentration added during sample preparation (10 mM, Methods), corresponding to an encapsulation efficiency of 100%. For each vesicle, a local lumen-dye density, *ρ*, was initially calculated as:

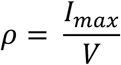

Where *I*_max_ is the maximum fluorescence response of the lumen-dye in arbitrary units and *V* is the vesicle volume in liters. The vesicle volume was calculated from the vesicle radius, which was determined based on the fluorescence intensity of the membrane-dye (see Methods, eq. (1)):

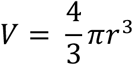

Where r is the vesicle radius. We considered the top 2% of the resulting lumen-dye densities as outliers and calculated the highest achievable density, πmax, as the mean of the top 2% of the remaining vesicles. All vesicles were subsequently calibrated into absolute lumen-dye concentration using the following equation:

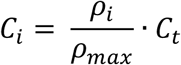

Where *C_i_* is the concentration of lumen-dye and *Ct* is the total concentration of dye added during sample preparation. See also Fig. S4.

## Supplementary discussion

### Physical model of calcium pumping and calcium leakage in a vesicle

We utilized an adapted form of the physical model that we developed in previous work to extract the pumping rates and permeabilities from single-molecule data^5,6,39^. The kinetics of calcium pumping into single vesicles by LMCA1 or hSERCA is described by the following system of equations:

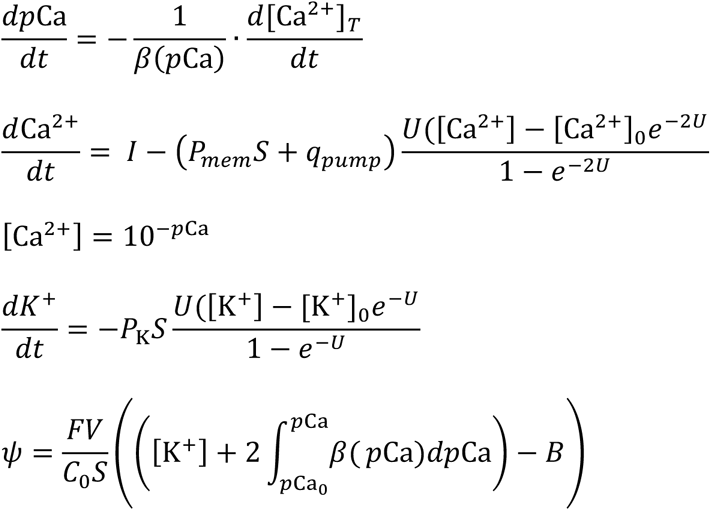

where U is the reduced membrane potential y/k_B_T, which is related to the net charge accumulated in the vesicle using a capacitor model for the membrane. We assume that the membrane potential outside the vesicle is zero. Bracketed quantities refer to molar quantities or concentrations, while unbracketed quantities are numbers. Hence, [X] = X/V, where V is the volume. Ca^2+^ is the total number of calcium ions that *enter* the lumen, and a fraction of these ions become buffered. Calcium pumping is represented by *I*, *P_mem_* is the passive calcium permeability of the membrane, *P*_pump_ is the calcium permeability associated with the pump that can also be expressed as *q*_pump_ (a transprotein calcium leak with units of permeability per unit area). The buffering capacity of the lumen (*b*) depends on the intracellular *p*Ca as discussed next. Both the volume (*V*) and the surface area (*S*) of the vesicle are assumed to remain fixed during the time course of calcium pumping. Quantities without subscripts refer to intracellular values, while a zero subscript ([…]_0_) refers to extracellular, fixed values. The integral in the equation for the membrane potential is the sum of all charged calcium ions that have entered the lumen during calcium pumping, Ca^2+^_T_. We set this value to zero at the beginning of all calculations, which essentially assumes that the luminal calcium ions initially do not significantly contribute to the resting membrane potential. Any trapped negative charges or partial negative charges, TRIS and Cl^--^, are modelled by a Donnan particle concentration (*B*), which is constant in time and set as described later. The K^+^ permeability (*P_K_*) is set to a large value since the presence of valinomycin in solution allows for fast shuttling of potassium across the bilayer. Please refer to table S1 for all values. Calcium pumping (*I*) is a time dependent quantity initiated at time t_on_ and then turned off at time t_off_, where both times are determined for each experimental trace prior to solving the system of equations. The initial [Ca^2+^] in the vesicle is set to either 1 µM or 0.5 µM depending on the experimental condition. The geometry of the system is assumed to be spherical with the vesicle radius (r) derived from measurements of the fluorescence intensity of the lipid-conjugated membrane marker according to eq. (1) for LMCA1, and directly from cryoEM size distributions for hSERCA (see Fig. S17). Once r is experimentally determined both *V* and *S* are also defined. The equations are solved with MATLAB using the *ode15s* stiff solver. The initial luminal potassium concentration and the value *B* in membrane potential are set so that the initial potassium flux and calcium flux across the membrane is zero; hence, the system starts at steady state.

### Simulating activity traces using the physical model

The physical model was adapted as described above and used to study the dependence of *p*Ca = -log[Ca^2+^] and the temporal fluorescence response, passive permeability, liposome radius and pumping rate. We utilized the model to simulate our experimental system with single LMCA1-facilitated calcium pumping into the lumen of single vesicles upon the addition of ATP. By varying the permeability in the model, whilst keeping all other parameters fixed (r = 30 nm, pumping rate = 1 ion/s), we found conditions where the resultant dynamic equilibrium was below dye saturation and exceeded the noise-level of the system to be able to continuously detect it (Fig. 1d).

To be able to relate these kinetic traces to the equivalent in fluorescence intensity recorded using the assay, we applied a transformation between [Ca^2+^] and fluorescence:

Here, we initially simulated pCa traces and then converted them into integrated fluorescence intensity using the experimentally determined calibration curve (FIG SX), and the following equation:

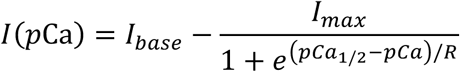

Where Imax is the maximum fluorescence, Ibase is the baseline fluorescence and *R* is the growth rate constant. This resulted in a normalized intensity trace with a baseline of 1. For r = 30 nm vesicles, the baseline intensity was scaled to 30 photons representing a typical experimental photon count (See Fig. S8). Due to the low typical photon count, we used an Electron Multiplying Charge-Coupled Device (EMCCD) as the preferred detector, hence applying both Poissionian noise and a multiplicative noise factor of 1.41 to the simulated intensity traces^55^.

### Signal intensity of single vesicles and choice of detector type

To determine the average photons emitted per vesicle, we imaged vesicles under laser illumination for 36 frames in buffer without ATP. We applied a ROI of 5×5 pixels and determined the mean integrated intensity per vesicle, which was converted into photons using the equation:

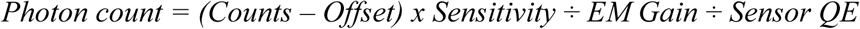

where *Counts* is the integrated intensity, *Offset* is the average dark signal (measured in the absence of laser illumination), *Sensitivity* is the sensitivity of the Andor iXon EM-CCD (using 17 MHz, 16-bit, the single EM amplifier was 5.05, *EM Gain* was 300 and we used a sensor *QE* of 87.5% at *λ* = 488 nm, using the camera quantum efficiency curves). The distribution was fitted using a lognormal distribution with an offset of 10, giving a mode of ∼29 photons (Fig. S8). Thus, we chose to perform our comparative simulations using 30 photons for the starting photon count per liposome.

The camera chosen for experiments depends on the amount of photon incidents on the detector. For low light levels (< 200 photons s^-1^), an EMCCD provides the optimum detector, benefitting from its high sensitivity towards low photon counts. For higher light levels (> 200 photons s^-1^), a Scientific Complementary Metal-oxide-Semiconductor image detector (sCMOS) becomes the better detector to use.

The number of photons that can be emitted from a vesicle depends on the number of encapsulated fluorophores. Since we determined experimentally that vesicles with r = 30 nm generated typically ∼30 photons s^-1^, we ran simulations for these vesicles assuming the properties of an EMCCD. Since the number of fluorophores encapsulated scales with vesicle volume, the number of photons emitted will scale with vesicle size. Thus, for vesicles larger than 30 nm (r = 100 nm and r = 1000 nm), we reach a regime where working with a sCMOS detector is beneficial over an EMCCD (Fig. S8). For simulations of large vesicles, the characteristics of a sCMOS detector were assumed and therefore, only Poissionian noise was imposed onto the intensity response. To avoid camera saturation, we restricted the camera exposure time, ensuring that the photon count did not exceed 2/3 of the full well capacity (15000 electrons using a 0.24 conversion factor).

### Defining the detection limits of the assay

We defined the upper detection limit of [Ca^2+^] of the assay to be 100 µM, where the dye response was assumed to be saturated. We determined experimentally that 100 µM Ca^2+^ provided ∼95% of the maximum fluorescence intensity of Cal-520, which could be achieved in excess of Ca^2+^ (2 mM) (see fig. 1b). In all simulations, the maximum resolvable pumping rate (I_max_) was thus defined as traces where [Ca^2+^]_lum_ exactly reached the saturation threshold of 100 µM. The minimum pumping rate (*I*_min_) that could be resolved was defined as when the maximum signal-to-noise (*SNR*) of the data was equal to a SNR threshold (SNR = 5).

Here, the *SNR* was defined as the ratio of the change in signal intensity to the noise of the initial baseline intensity, SNR = (*I*_norm,max_ - 1)/*σ*_norm,base_). The SNR was determined by applying exponential fits to the intensity response and determining the maximum value.

### Simulating the detection range as a function of vesicle permeability, size and experimental time window

We utilized the model to simulate our experimental system with an initial calcium concentration [Ca]_start_ = 0.5 µM, and after 5 minutes, a buffer containing 1 mM ATP was added to initiate Ca^2+^ pumping and we then monitored the fluorescence intensity response for 35 minutes. This allowed us to determine which permeability would generate a fluorescence intensity response corresponding to the upper- and lower detection limit of the system (*I*_max_, *I*_min_) for a given pumping rate. The resulting plots (Fig. 1d and Fig. S8) were generated by running two different simulations to determine the pumping rate at which the SNR threshold and saturation threshold was met, respectively. For the former, we started by choosing a low pumping rate and a permeability value and then used the model to simulate a fluorescence intensity trace including noise. We then fitted the trace using an exponential fit to recover the maximum SNR that was achieved within the experimental window. If the SNR was less than the threshold, we increased the pumping rate and repeated the simulation until the SNR threshold was met.

To determine the pumping rate for which the intensity response would saturate for a given permeability, we used a similar approach. However, instead of applying the transformation from [Ca^2+^] to fluorescence intensity and applying noise, we simply checked whether the [Ca^2+^]_lum_ exceeded the saturation threshold within the given experimental window.

In Fig. S10, we used simulations to investigate the detection range of the assay as a function of size. Here, we ran simulations for three different vesicle sizes (r=30 nm, r=100 nm and r=1000 nm). To determine the pumping rate for which the SNR threshold was met, and thus the lowest detectable current (*I*_min_), we fixed the permeability to a low value (*P*_mem_ = 10^-10^ cm s^-1^), where passive calcium leakage could be considered insignificant. The saturation threshold (*I*_max_) was determined by assuming a permeability of *P*_mem_ = 10^-6^ cm s^-1^.

In Fig. S8, we simulated activity traces with experimental time-windows of 5 mins, 35 mins and 2 hours and plotted the resulting *I*_min_ as a function of *P*_mem_ for all conditions. By extending the experimental time-window from 5 mins to 2 hours, *I*_min_ is reduced from ∼0.05 ions s^-1^ (∼16 zA) to ∼0.0025 ions s^-1^ (0.8 zA) in vesicles with low permeability (*P*_mem_ ≲ 10^-7^ cm s^-1^), where the duration of the measurement is limiting the build-up of [Ca^2+^]_lum_.

In addition to increasing the experimental time-window, the sensitivity of the assay can, in principle, be improved by implementing a fluorescent indicator with lower *K*_d_ or achieving higher fluorescent signal by having increased photostability or encapsulation efficiency of the indicator.

### Influence of pumping rate, permeability and vesicle size on SNR and response time of the assay

In Fig. S11d, we simulated three activity traces with fixed permeabilities and sizes (*P*_mem_ = 10^-6^ cm s^-1^, r = 30 nm), while varying the pumping rate from 0.1 to 10 ions ^-1^ (32 to 3200 zA). We then normalized the traces to the fluorescence intensity at the baseline (a) or their respective dynamic equilibrium (b). We observe that with increasing pumping rate, the amplitude (and therefore the SNR) of the response increases and the response times becomes faster. Hence, high pumping rates are more easily resolved by the assay, compared to slow pumping rates, because they provide faster and brighter transporter-mediated responses.

In Fig. S10 c-d, we simulated three activity traces with fixed pumping rates and sizes (I = 0.05 ions s-1 (16 zA), r = 30 nm), while varying the permeability from 10^-6^ to 10^-7^ cm s^-1^. We observe that with increasing permeability, the SNR of the response decreases while the response times becomes faster. Hence, while permeability can be increased to speed up the response times of the assay, the SNR of the response is sacrificed correspondingly.

In Fig. S10 e-f, we simulated three activity traces with fixed pumping rates and permeabilities (I = 0.05 ions s-1 (16 zA), P_mem_ = 10^-6^ cm s^-1^), while varying the vesicle radius from 30 to 50 nm. We observe that with increasing size, the SNR of the response decreases and the response times becomes slower. Hence, small vesicle sizes provides both a high sensitivity and fast response times compared to larger vesicles.

We defined the response time to transporter-mediated Ca^2+^ influx (τ_influx_) as the time it takes for the active pump to build up 1-e^-1^ (∼63%) of the intensity at dynamic equilibrium and the response time to passive Ca^2+^ efflux during resting periods of the pump as the time it takes for the intensity to decay to e^-1^ (∼37%) (Fig. 1f and Fig. S9).

We found that τ_efflux_ was slower than τ_influx_ for any given permeability, hence we used τ_efflux_ as a measure of the minimum detectable dwell-times (*t*_*min*_) in all simulations (Fig. S9). However, while low permeability (*P*_mem_ ≤ 10^-6^ cm s^-1^) defines a regime where influx kinetics are much faster than efflux kinetics (τ_influx_ ≪ τ_efflux_), at high permeability (*P*_mem_ ≥ 10^-5^ cm s^-1^), influx and efflux kinetics approach similar values (*t*_influx_ ≈ *t*_efflux_) (Fig. S9).

As discussed previously, the assay resolves the typical pumping rates of LMCA1 (*I* ≈1 s^-1^) as well as active/inactive mode dwell-times (t_min_ ≥0.5 min). However, if the pumping rate is sufficiently high (I ≫1 s^-1^), it is in principle possible for the assay to resolve much faster mode-switching (t_min_ ≪ 0.5 min) since the permeability can be increased to compensate for the faster pumping rate (Fig. S11). Equally, the restriction of working with small vesicles (r ≈ 30 nm) becomes less important when resolving fast pumping rates, as shown in Fig. S11, where vesicles with r = 50 nm or r = 100 nm can be used to resolve a pumping rate of I = 100 s^-1^ and dwell-times down to t_min_ ≈ 10^-2^ min.

## Supplementary figures

**Figure S1.**
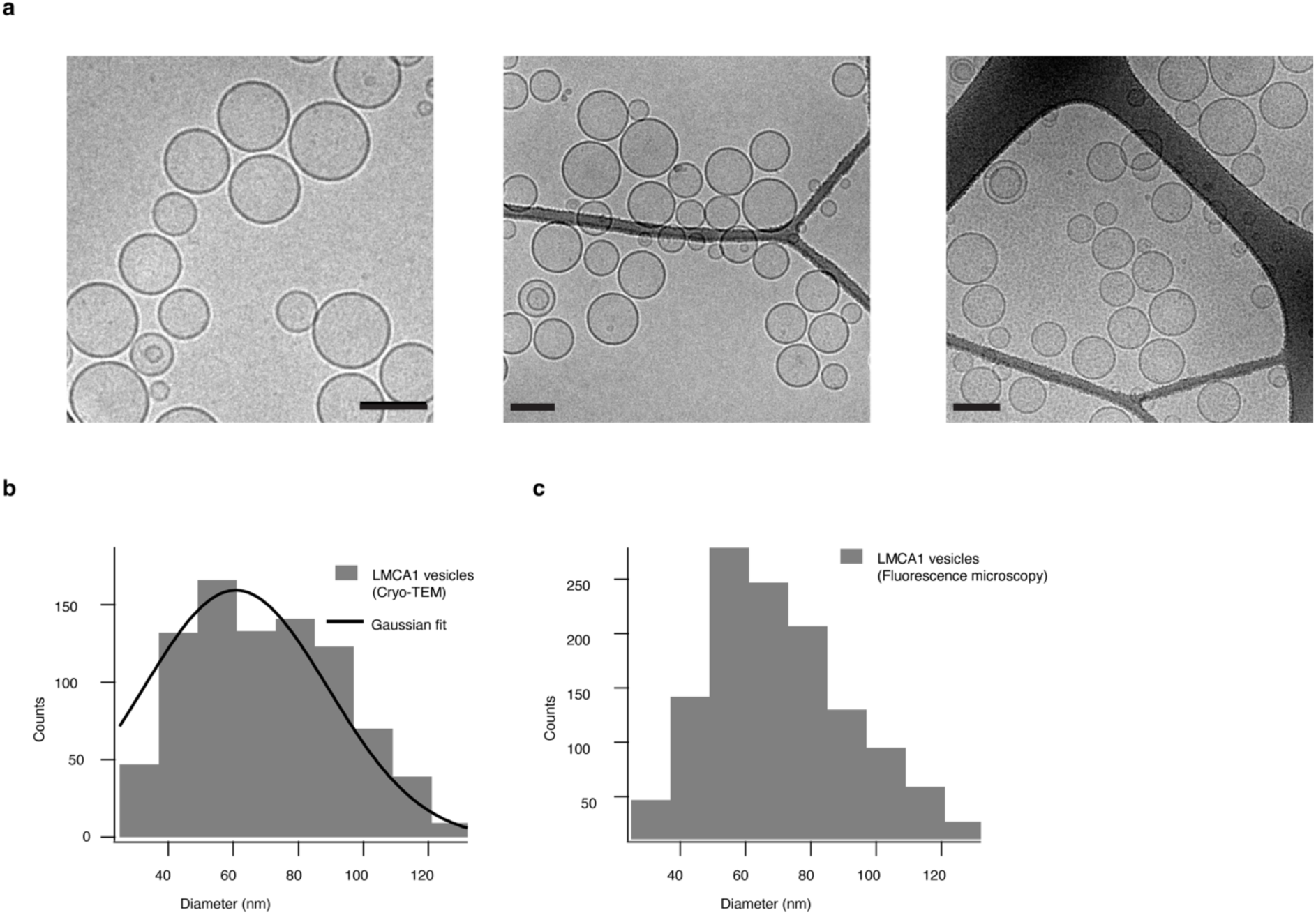
Size and morphology of LMCA1 reconstituted proteoliposomes (**a**) Representative micrographs of LMCA1-reconstituted vesicles. The vesicles were predominantly unilamellar and spherical. All scale bars are 100 nm. (**b**) Vesicle size distribution as measured by Cryo-TEM, fitted with a Gaussian function with a center of 60 ± 2 nm (mean ± HWHM). (**c**) Size distributions of the vesicles as determined by wide-field fluorescence microscopy (see Methods). The histograms in (**b**) and (**c**) were based on *n*=871 and *n*=1394 vesicles, respectively.

**Fig S2.**
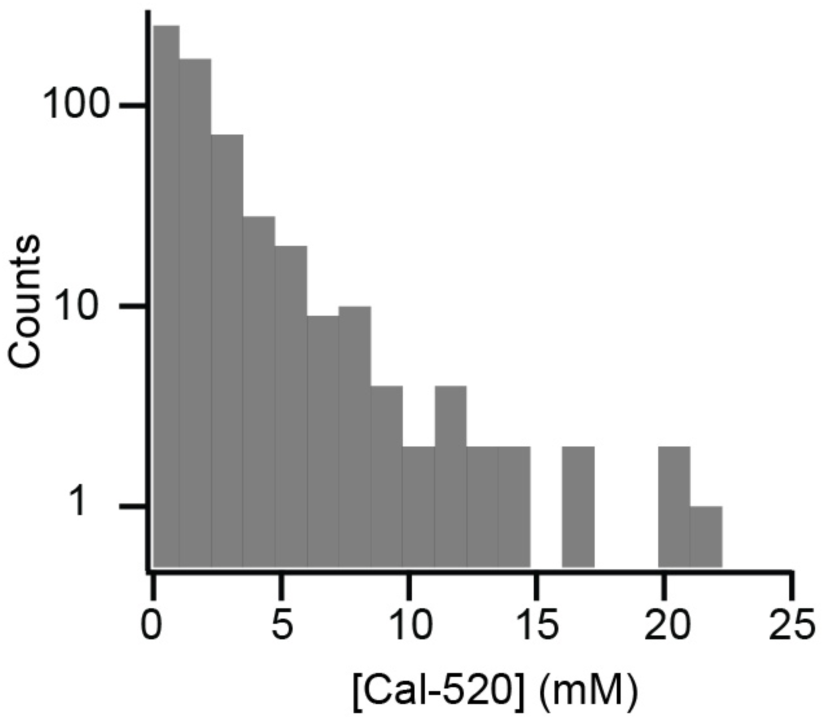
Local determination of lumen-dye concentration reveals mean luminal concentration of ∼2 mM. (a) Distribution of lumen-dye concentration ([Cal-520]) in a sample of loaded vesicles (see also supplementary methods). *n* = 583 single vesicles. Empty vesicles are not included in the histogram.

**Figure S3.**
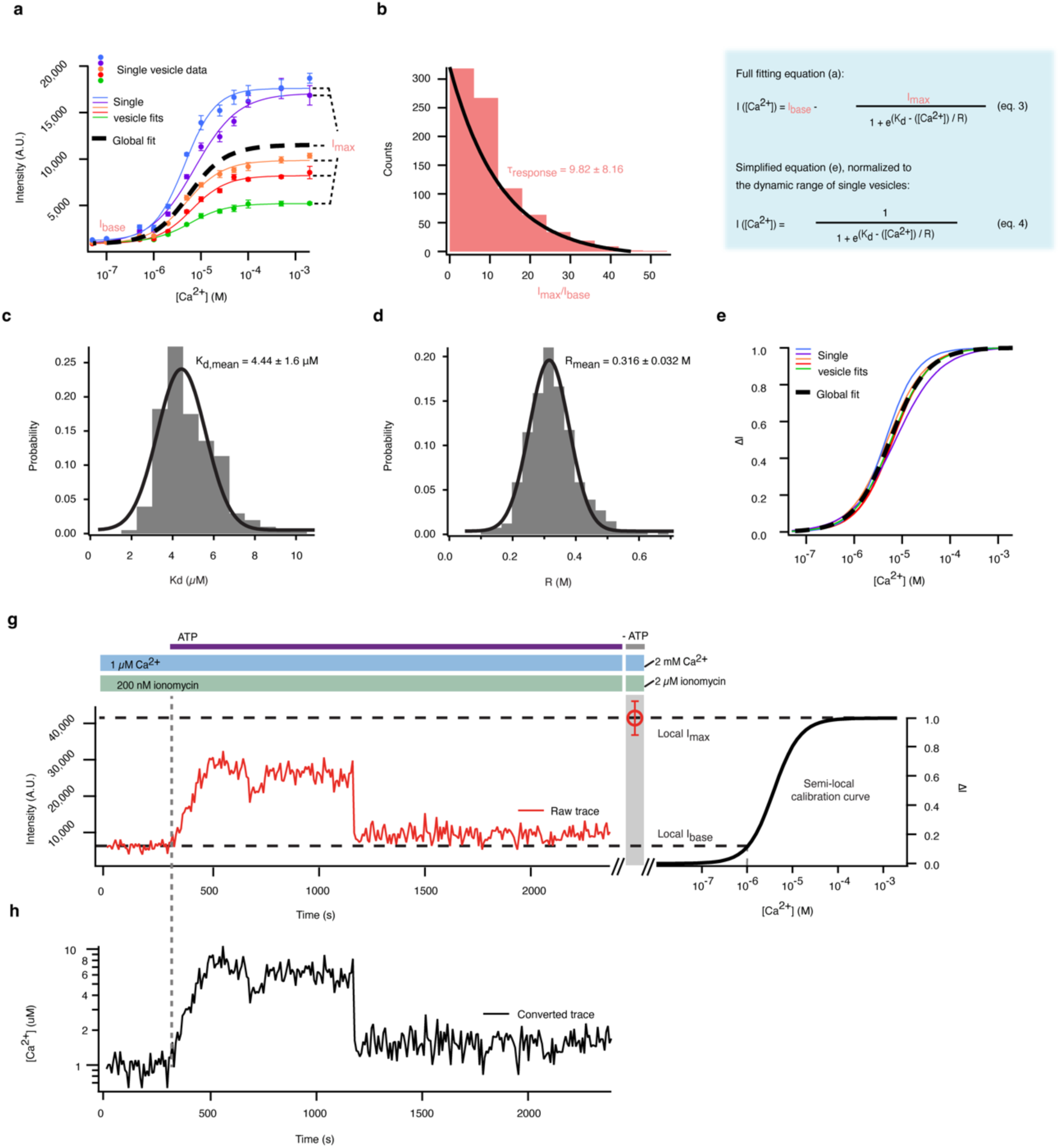
Method for conversion of kinetic traces from fluorescence intensity into Ca^2+^ concentration depends on local determination of dynamic range. **a**, We titrated vesicles with Ca^2+^ calibration buffers and monitored the fluorescence intensity of the Ca^2+^ indicator as a function of [Ca^2+^]. Five representative single-vesicle dose-response curves fitted to sigmoidal functions (solid-coloured traces). The curves displayed a high degree of vesicle-to-vesicle heterogeneity. Data points are mean ± s.d. for *n* = 3 consecutively recorded images of the same single vesicle. Global fit (dashed black lines) represents the mean of the displayed single-vesicle fits. **b-d**, Distributions of Sigmoidal fit parameters indicated that the vesicle-to-vesicle heterogeneity observed in (**a**) was vastly dominated by a large spread in the dynamic range (*I*_max_/*I*_base_) (**b**), while the dissociation constant, *K*_d_ (**c**), and growth rate, *R* (**d**), were characterized by narrow normal distributions with mean values of *K*_d,mean_ = 4.4 ± 1.6 µM and *R*_mean_ = 0.316 ± 0.032 M. **e**, Shows the same single-vesicle fits as in (**a**), but normalized to the dynamic range of each vesicle (ΔI). Vesicle-to-vesicle heterogeneity is reduced in (**e**) compared to (**a**), because it only reflects variations in *K*_d_ and *R*, and not the dynamic range. To overcome the challenge of vesicle-to-vesicle heterogeneity in dynamic range, we experimentally determined *I*_base_ and *I_max_* at the single-vesicle level pre-and post all activity measurements (**g**). We combined local values (*I*_max_ and *I*_base_) with global values (*K*_d,mean_ and *R*_mean_) to generate semi-local calibration curves that we could use to map normalized intensity traces onto the corresponding [Ca^2+^] values, as shown in (**g-h**).

**Figure S4.**
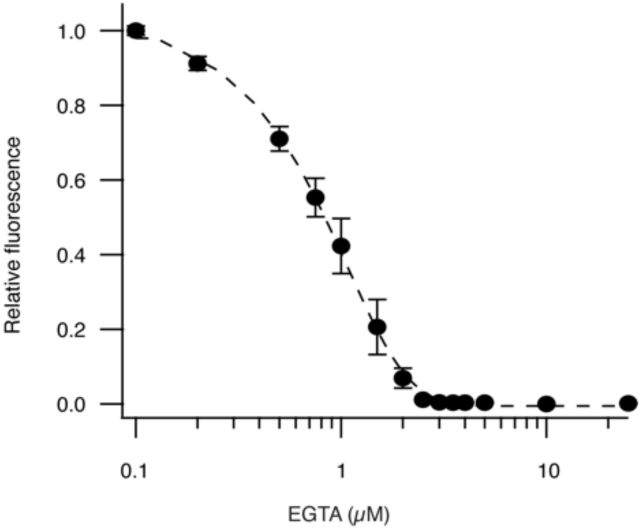
Reducing background fluorescence by chelation of Ca^2+^ contaminants. (**a**) A buffer solution (see Supplementary methods) containing Ca^2+^ indicator Fluo-5N was prepared from Milli-Q water and titrated with EGTA while monitoring the fluorescence intensity using a spectrofluorometer. The intensity of the Ca^2+^ indicator decreased as EGTA chelated free Ca^2+^ ions in the buffer and reached a plateau at ∼3 μM EGTA, indicating that all Ca^2+^ ions were chelated. Thus, all buffers used during this study were initially added 3 μM EGTA, unless otherwise stated, to reduce background fluorescence and accurately control [Ca^2+^]. Data points are mean ± s.d. for *n* = 5.

**Figure S5.**
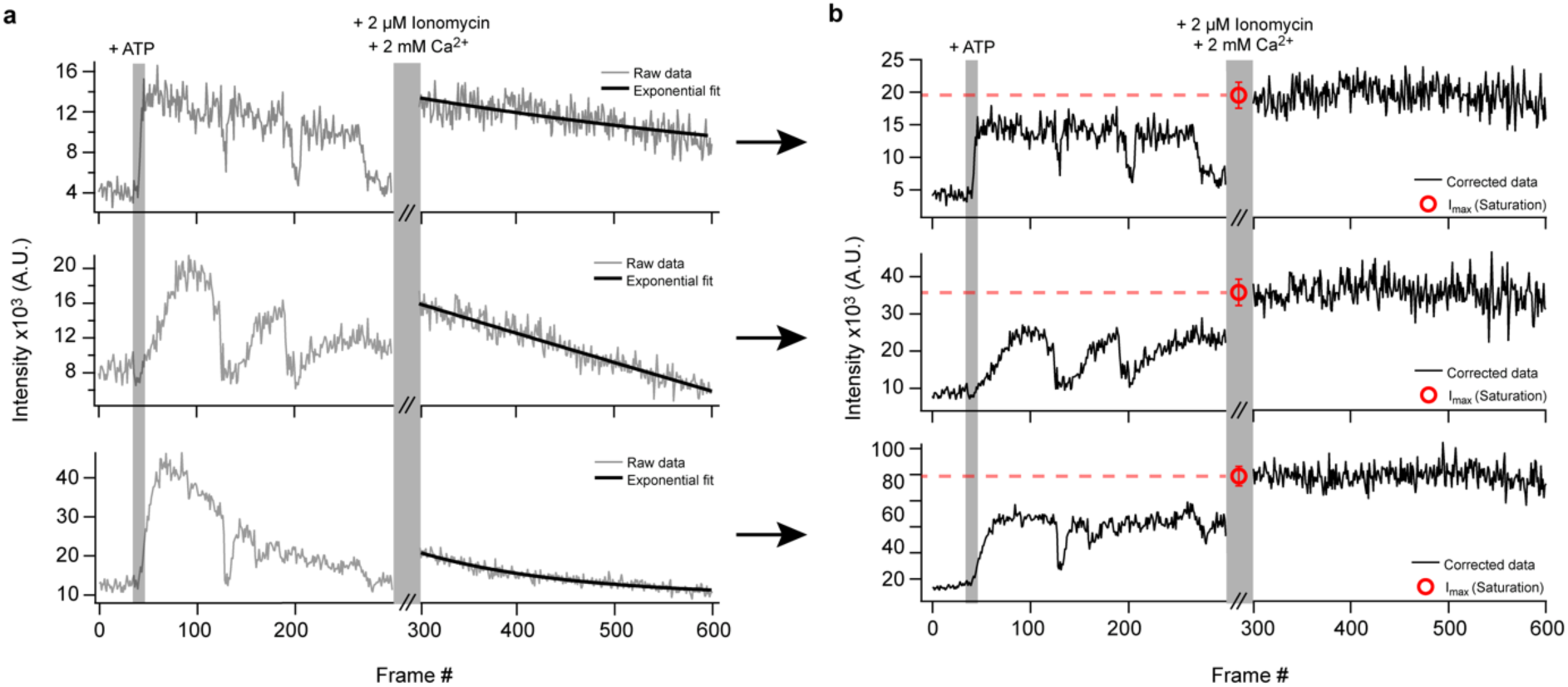
Local correction for photobleaching compensates for a continuous signal decay. (**a**) Three typical LMCA1 single-molecule activity traces (frame # 0-300) and their corresponding photobleaching measurements (frame # 301-600). LMCA1 activity was initiated by the addition of 1 mM ATP. After 300 frames, the image acquisition was stopped, and the vesicles were incubated for 10 minutes with saturation buffer including 2 mM Ca^2+^ and 2 µM ionomycin to collapse any established Ca^2+^ gradients and saturate the Ca^2+^ indicator. The vesicles were subsequently imaged for another 300 frames to characterize the local photobleaching kinetics. Photobleaching traces were fitted with an exponential decay function which was subsequently used for correcting the entire image sequence for photobleaching-induced signal decay. **b**, The three traces from (a) after local correction for photobleaching. The saturation intensity, *I*_max_, was calculated as the mean of the initial 50 frames of the corrected photobleaching trace and used for semi-local calibration from intensity to [Ca^2+^].

**Figure S6.**
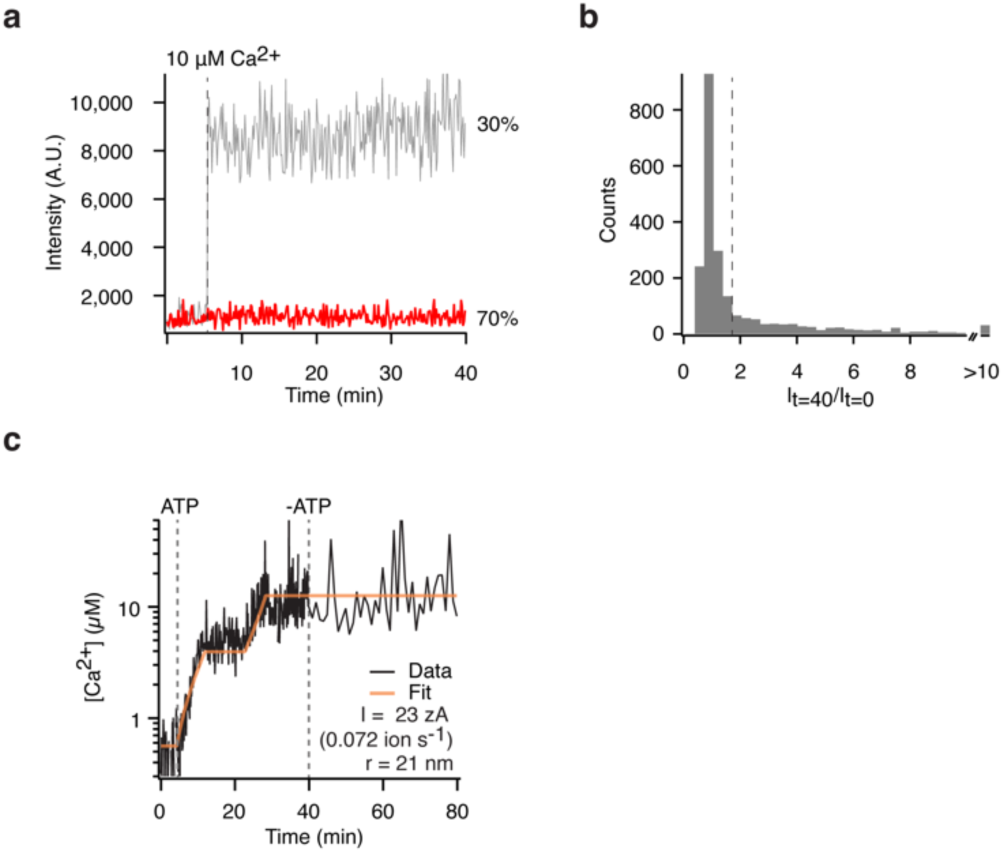
Intact vesicles are impermeable to Ca^2+^ for the duration of a typical experiment and demonstrate LMCA1 mediated Ca^2+^ accumulation in the lumen. **a**, Representative kinetics of Ca^2+^ influx in single vesicles in response to an imposed 10 μM Ca^2+^ gradient. Most vesicles were intact and did not display any detectable leakage of Ca^2+^ over a 35-minute measurement (red trace, “tight”), while a subpopulation of vesicles leaked “immediately” within a single frame (grey trace, “leaky”). **b**, Histogram of relative fluorescence response of single vesicles as measured before (*I*_t=0_) and 35 min after (*I*_t=40_) injection of 10 μM Ca^2+^ showing that 70% of the vesicles are tight (*I*_t=40_/*I*_t=0_ < 1.55). Data are from 2201 single vesicles. *n* = 3 replicates. **c,** Example LMCA1-mediated Ca^2+^ pumping kinetics in a single vesicle upon addition and subsequent removal of ATP. For vesicles impermeable to Ca^2+^ switching of LMCA1 results in a step-like increases in fluorescence. Kinetics are fitted to a quantitative model (orange trace) to extract the pumping rate of 23 zA.

**Figure S7.**
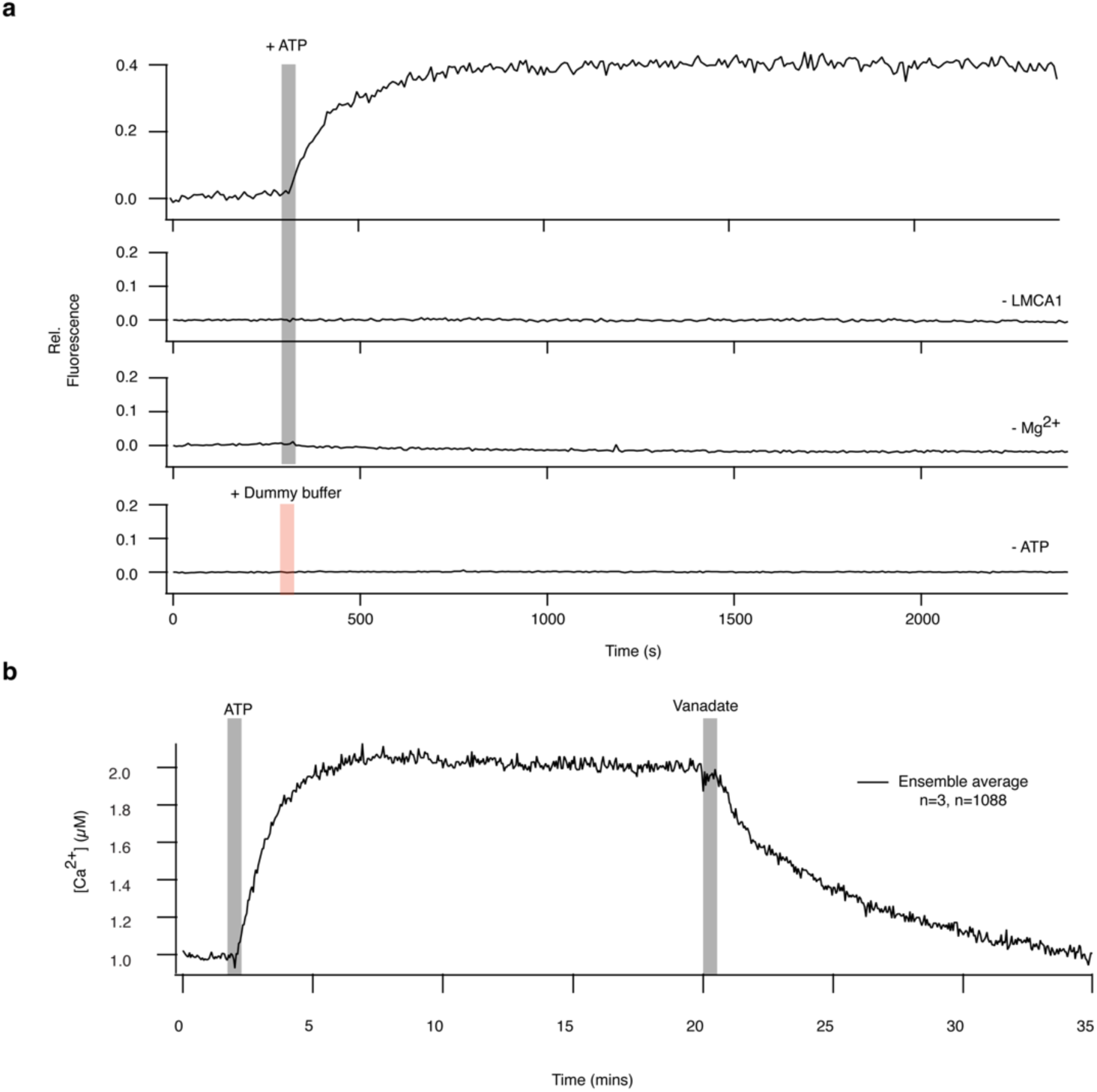
Ensemble average experiments confirm that activity is initiated by a calcium pumping ATPase and is inhibited by vanadate. **a**, Positive and negative controls of ensemble-averaged pumping kinetics. Active Ca^2+^ transport was observed only in simultaneous presence of LMCA1, 1 mM ATP and 1 mM Mg^2+^. Data are mean of *n* > 400 single vesicles. *n*=3 replicates**. b,** All activity was blocked upon addition of the P-type ATPase specific inhibitor, vanadate.

**Figure S8.**
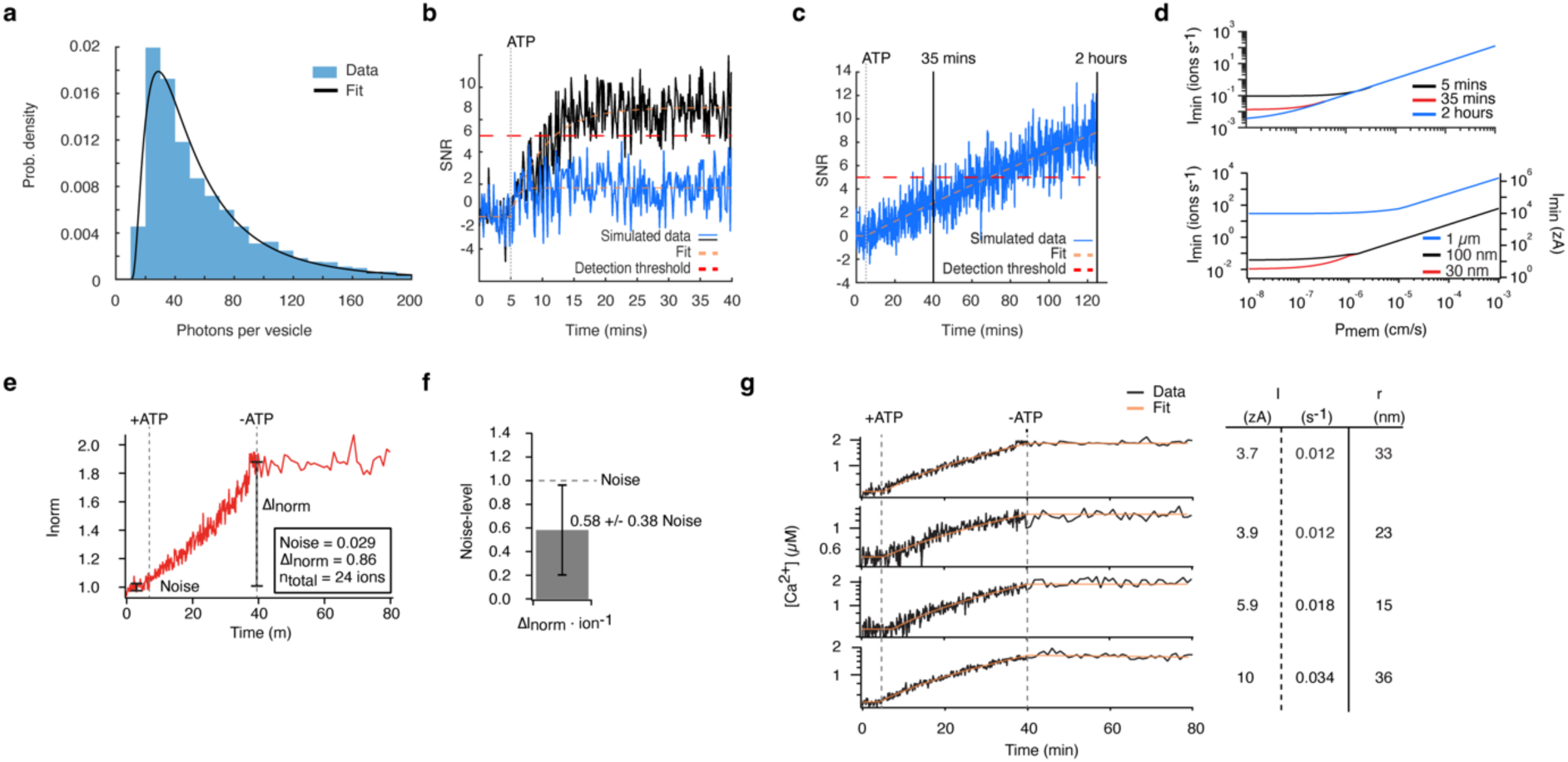
Simulations and experiments confirm the ability of the assay to measure zepto-Ampere currents by single LMCA1. The lowest detectable pumping rate of the assay is determined by the signal-to-noise, vesicle permeability, vesicle size and the duration of the experiment. **a**, Experimentally determined number of photons emitted per vesicle per frame in the transporter assay (see also supplementary methods). The distribution is fitted with a lognormal function with an offset of 10 giving a mode of 28.6 photons. *n* = 4635. The photon count per vesicle determines the noise-level of the measurements. **b**, Simulated activity traces with fixed pumping rate (I = 0.1 ions s^-^1 or 32 zA) and varying *P*_mem_ exemplifying a detectable (black trace) and a non-detectable trace (blue trace) within a 35 mins activity recording. The simulated data are fitted with exponential functions (dashed lines) to determine whether the detection threshold of SNR=5 is exceeded (red dashed line). **c**, Simulated activity trace of a slow pumping rate (blue trace, 0.0075 ions s^-1^, 2.4 zA) during a 2-hour activity recording exemplifying that by extending the duration of the me experiment from 35 mins to 2 hours, the assay can detect slower pumping rates which would otherwise have been considered below the detection limit. **d**, Top: Simulation of the lowest detectable pumping rate (*I*_min_) as a function of *P*_mem_ for 3 different experimental durations (black trace: 5 mins, red trace: 35 mins, blue trace: 2 hours), demonstrating that the assay can resolve lower pumping rates when the duration of the experiment is extended. Bottom: Simulation of *I*_min_ as a function of *P*_mem_ for 3 vesicle radii (red trace: 30nm, black trace: 100 nm, blue trace: 1 µm), demonstrating that small vesicles provide the ability resolve slower pumping rates compared to larger vesicles. **e,** Example of a normalized fluorescence response to ultra-slow Ca^2+^ pumping kinetics in a low-permeability vesicle. We determined the total fluorescence response achieved over 35 min of Ca^2+^ pumping (*ΔI*_norm_) and the noise-level of the kinetic recording as the std of the baseline (Noise-level), prior to the injection of ATP. The total number of transported ions (n_total_) were extracted using the physical model (Supplementary discussion). **f**, Mean fluorescence response per transported ion (ΔI_norm_ × ion^-1^), normalized to the noise-level of the kinetic recording. The fluorescence response generated by a single-ion translocation event was below the noise-level of the kinetic traces (0.58 ± 0.38) and could therefore not be directly resolved as discrete steps in fluorescence intensity. The analysis was based on *n* = 5 kinetic traces, displaying Ca^2+^ currents ≤ 10 zA. **g,** Representative LMCA1 Ca^2+^ pumping kinetics upon addition of ATP. The luminal [Ca^2+^] remained stable after removal of ATP, indicating that the vesicles had very low permeability to Ca^2+^. Vesicle sizes (*r*) were determined as described in the methods section. The data were fitted to a physical model of Ca^2+^ upconcentration in vesicles^2^ (orange trace) to extract single-molecule pumping rates (*I*).

**Figure S9.**
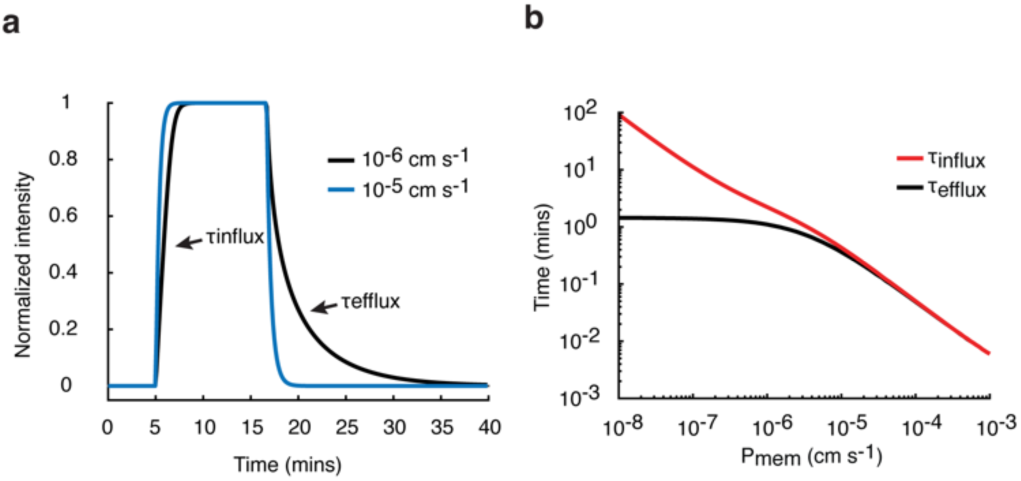
The kinetics of Ca^2+^ efflux during resting periods of the pump determines the response-time of the assay. **a**, Simulated activity traces with mode-switching events between active and inactive modes in vesicles with varied permeabilities. At low permeability (P_mem_ = 10^-6^ cm s^-1^, black trace), the Ca^2+^ efflux kinetics (τ_efflux_) is much slower than the influx kinetics (τ_influx_), while at high permeability (P_mem_ = 10^-5^ cm s^-1^, blue trace) the kinetics of τ_efflux_ and τ_influx_ are similar. **b**, τ_influx_ and τ_efflux_ as a function of P_mem_. The kinetics of τ_efflux_ is slower than τ_influx_ for any given permeability, hence τ_efflux_ provides the measure of the minimum detectable dwell-times (*t*_*min*_) of the assay.

**Figure S10.**
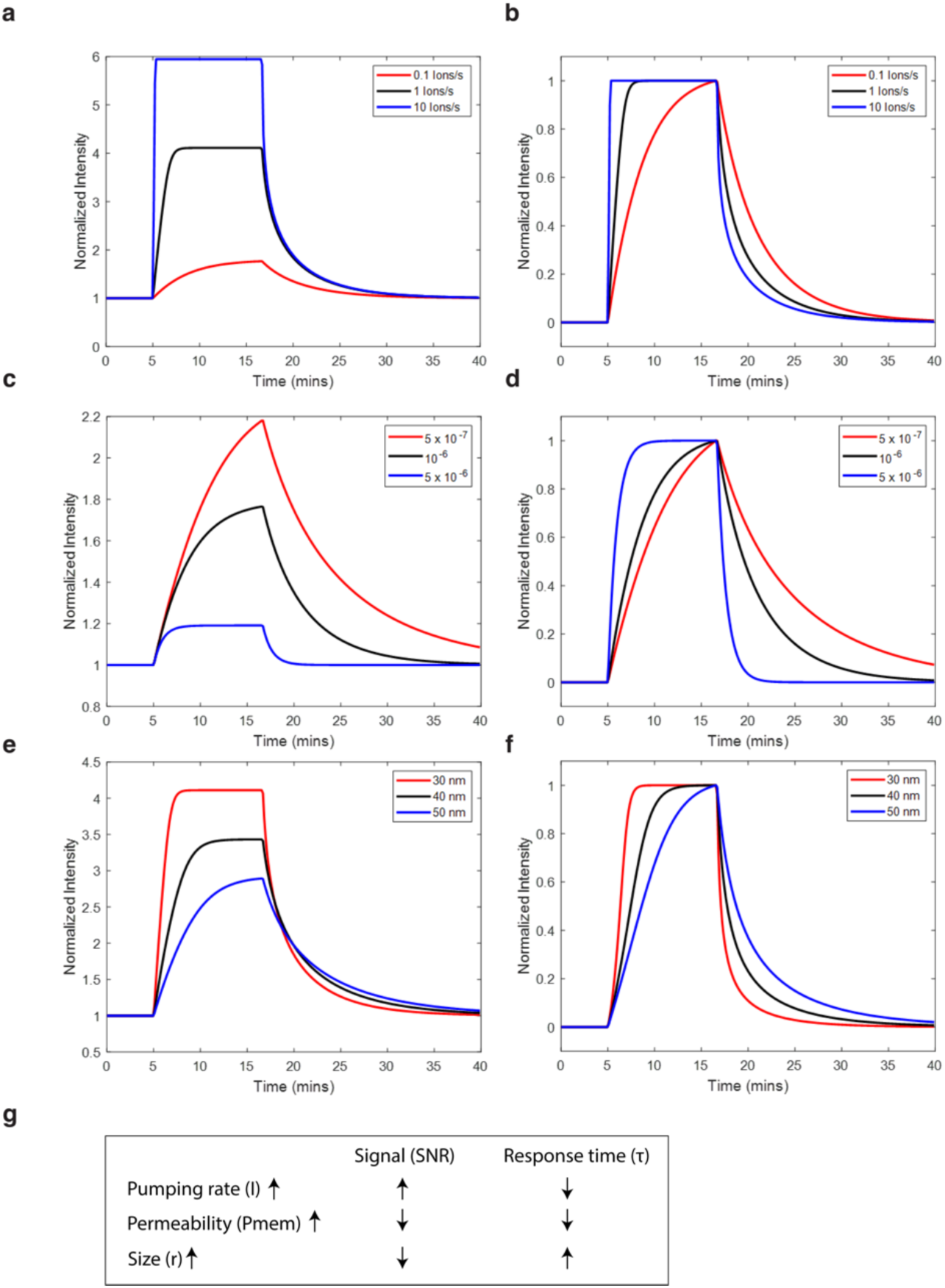
Influence of pumping rate, permeability and vesicle size on SNR and response time. Simulated activity traces with varying pumping rates **(a-b)**, vesicle permeability **(c-d)** or vesicle size **(e-f)**. Traces are normalized either to the baseline intensity (a, c, e) to indicate the the signal-to-noise ratio (SNR) of the response or to the intensity at the established dynamic equilibrium (b, d, f) to indicate the response time. High SNR can be achieved by high pumping rates (a), low vesicle permeability (c) and small vesicle sizes (e). Fast response times can be achieved by high pumping rates (b), high vesicle permeability (d) and small vesicle sizes (f). The effect of increasing the three parameters on SNR and response time is summarized in **g**.

**Figure S11.**
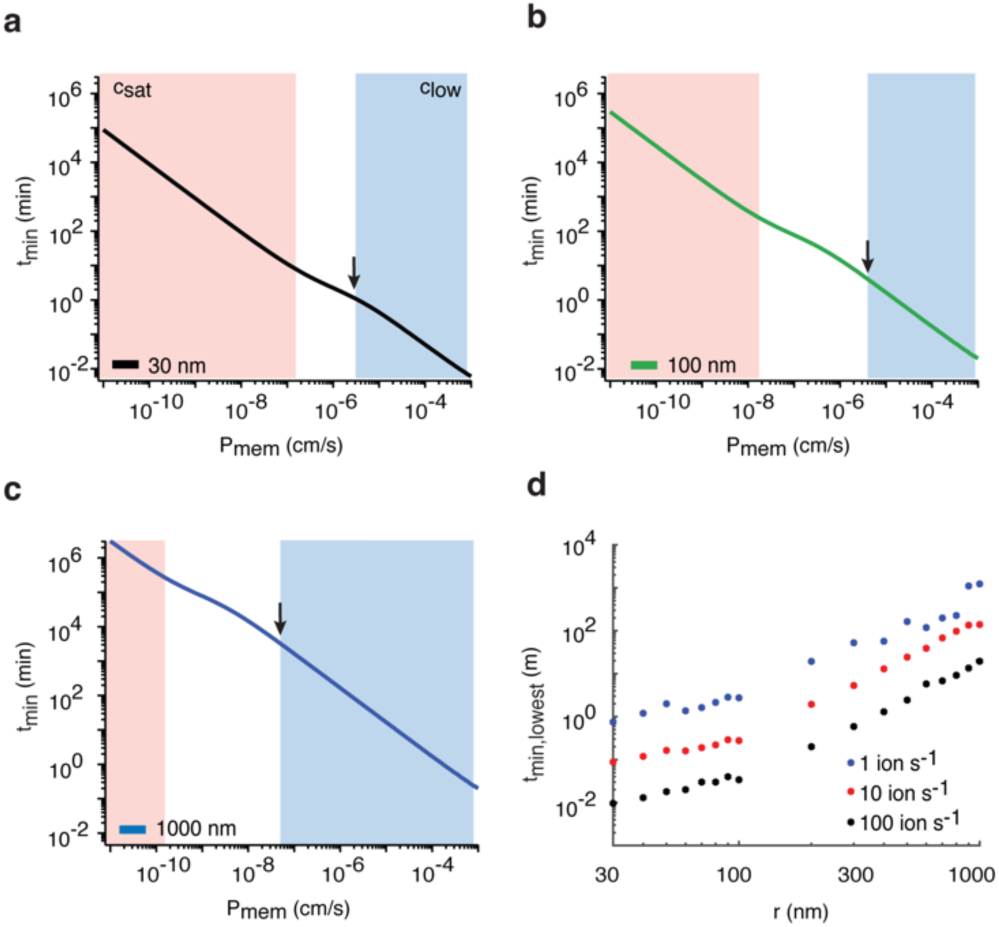
Small vesicle sizes are required for detecting mode-switching of slow-pumping transporter LMCA1. **a-c**, Minimum detectable dwell-times (t_min_) as function of P_mem_ for a typical LMCA1 pumping rate (I = 1 ion s^-1^) in a vesicle with a radius of r = 30 nm (**a**, black trace), r = 100 nm (**b**, green trace) or r = 1000 nm (**c**, green trace). t_min_ increases with increasing vesicle size thus resulting in slower response times of the assay. Arrows indicate how the lowest detectable dwell-time (t_min,lowest_) is determined for different vesicle sizes. **d**, t_min,lowest_ as function of r for a pumping rate of I = 1 ion s^-1^ (blue trace), I = 10 ions s^-1^ (red trace) or I = 100 ions s^-1^ (black trace). The requirement for using small vesicles (r=30nm) does not apply to transporters with high pumping rates (I = 10 ions s^-1^ and I = 100 ions s^-1^), for which t_min,lowest_ ≪ 1 minute when using a vesicle size of r ∼ 100nm.

**Figure S12.**
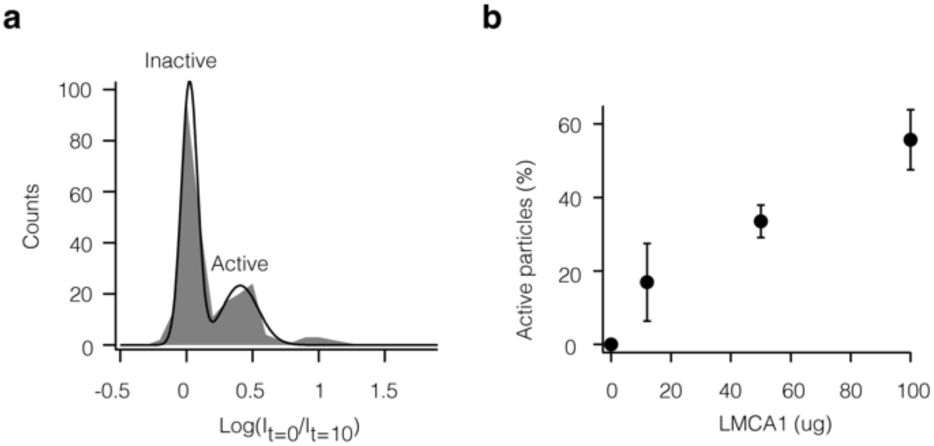
Dilution of the protein to lipid ratio ensures single copy numbers of LMCA1 per vesicle. (**a**) 68 ± 4% of all vesicles are inactive, indicating a low copy number of LMCA1 in the active vesicles. Distributions of relative fluorescence increases of single vesicles reconstituted LMCA1 measured before (*I_t=0_*) and after (*I_t=10_*) 10 min of incubation with 1 mM ATP. The fraction of vesicles containing at least one active transporter was determined as the relative area under the two peaks of a double Gaussian fit, indicating the active-(32 ± 4%) and inactive (68 ± 4%) fraction of the sample. (**b**) Titration of the protein to lipid ratio by adjusting the mass of reconstituted LMCA1 shows regulation of the fraction of active vesicles.

**Figure S13.**
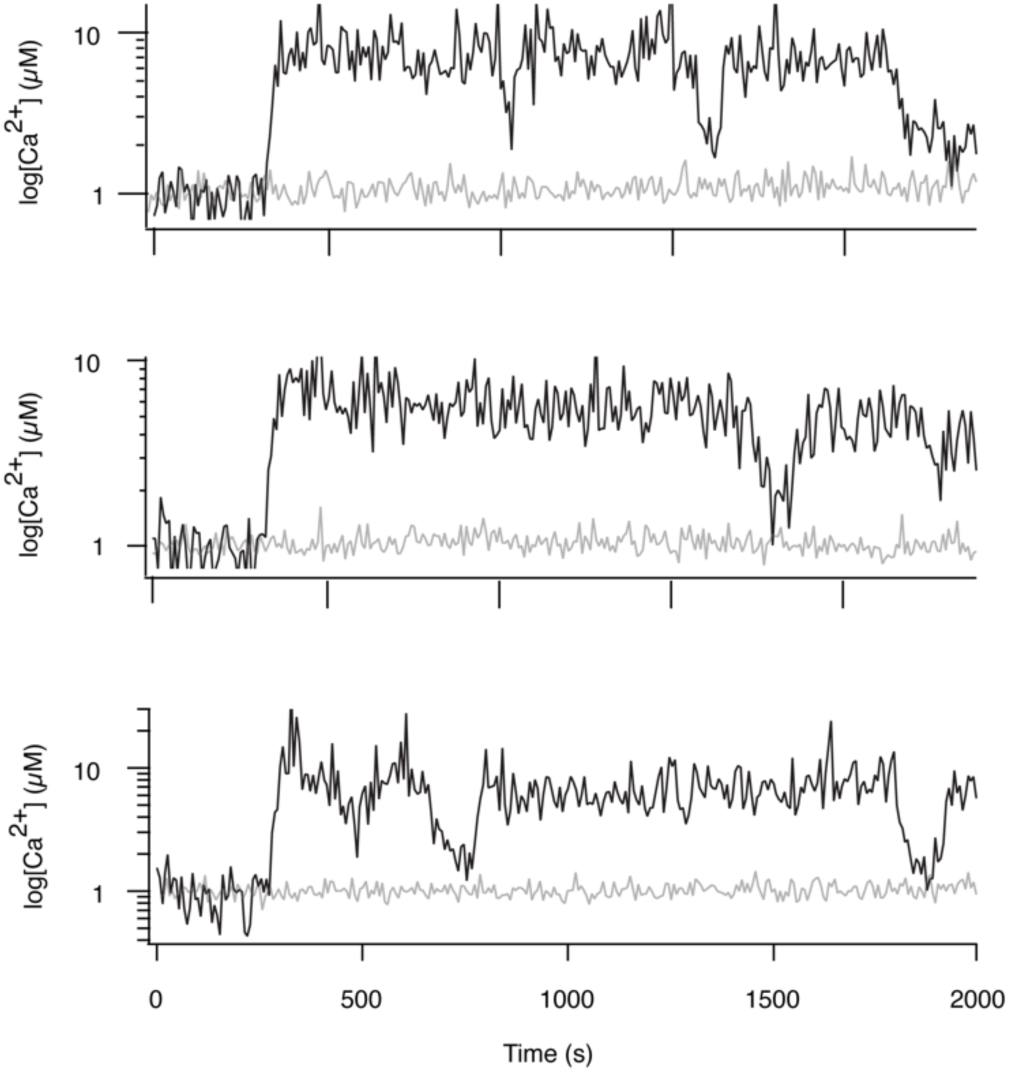
Additional long-term activity recordings of single LMCA1 pumps in single vesicles. Representative activity recordings of single LMCA1 pumps displaying stochastic switching between long-lived modes. Activity was initiated upon addition of 1mM ATP. Data are represented as the relative change in fluorescence response of the encapsulated Ca^2+^ indicator (Δ*I*).

**Fig S14.**
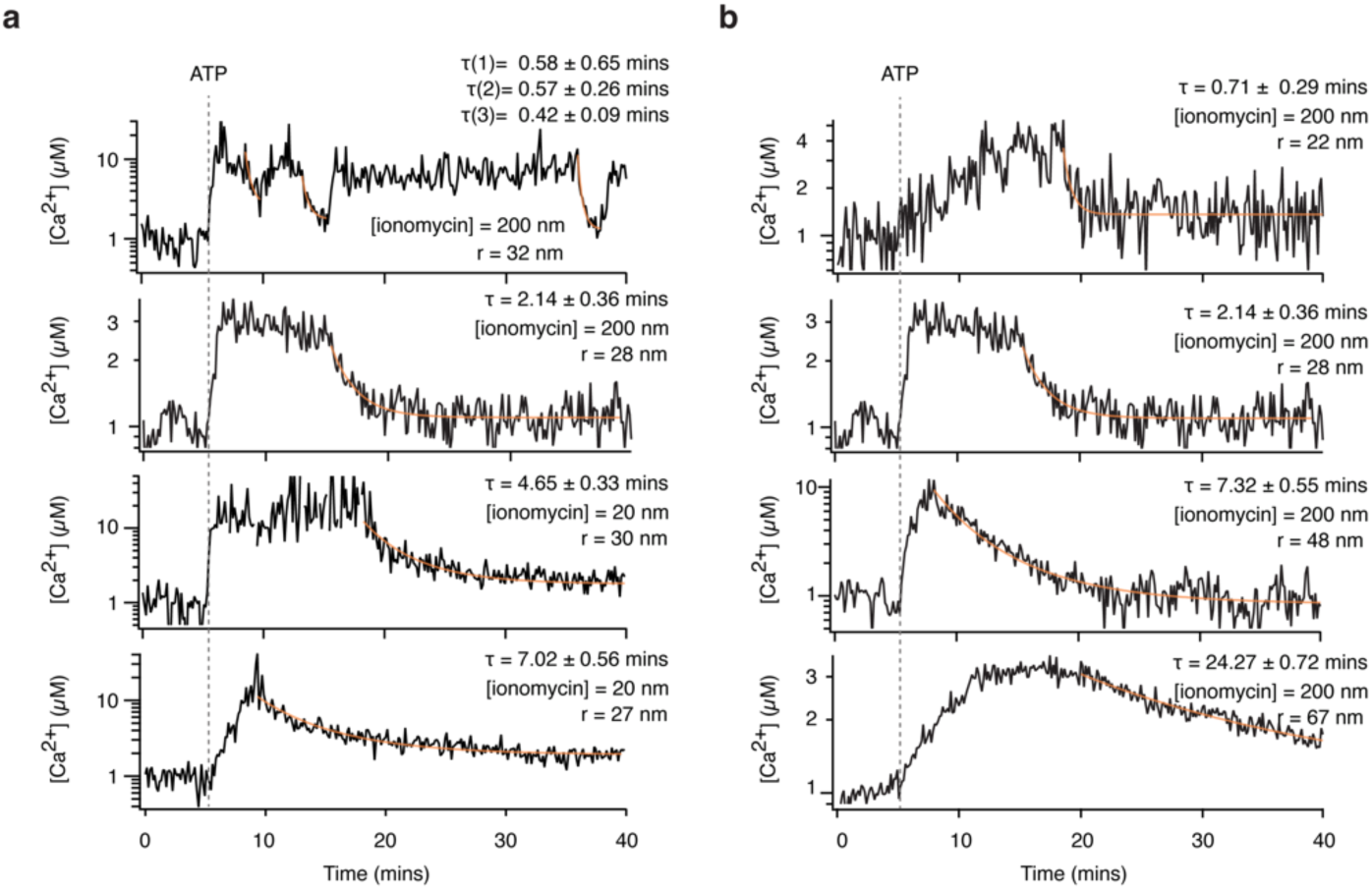
Additional controls: experimental validation of the dependence of response-time on membrane permeability and vesicle size. **a-b**, Representative single vesicle recordings of Ca^2+^ pumping and mode-switching in vesicles with varying membrane permeability (P_mem_) (**a**) or size (r) (**b**). With increasing concentration of ionomycin (*C*_iono_) we observed faster response times (τ_r_) in vesicles of similar sizes (r∼30 nm) (**a**), determined as the decay constant from an exponential fit to the decay in [Ca^2+^] during resting periods of LMCA1 (orange trace). (**b**) Increasing the vesicle size, without changing P_mem_ (*C*_iono_ = 200 nM), resulted in slower response times.

**Fig S15.**
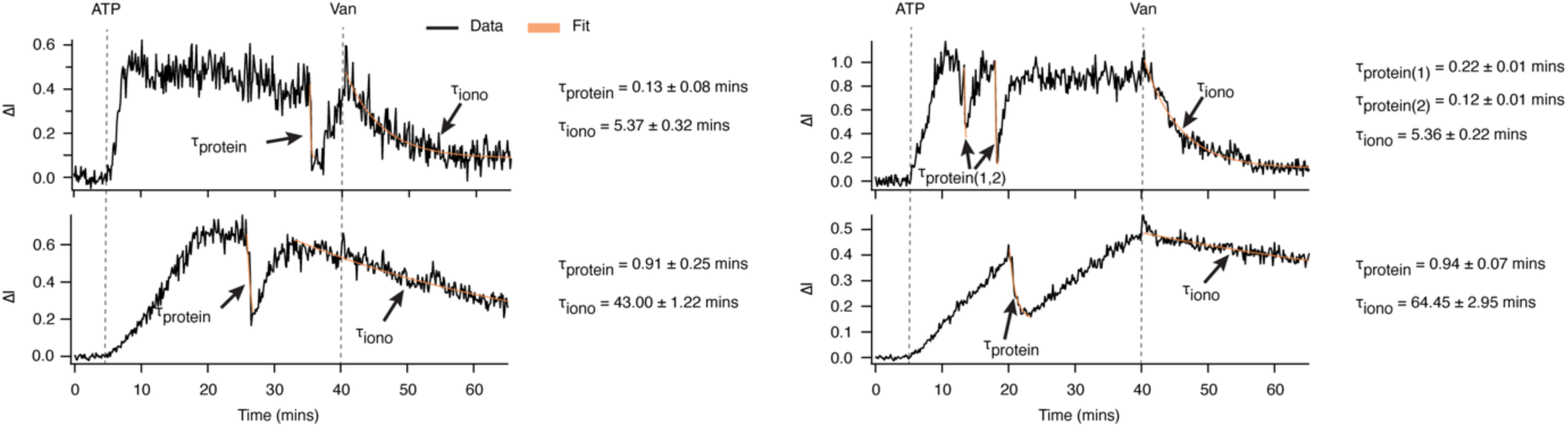
Additional single molecule traces displaying LMCA1 leakage currents. Representative single-vesicle recordings of Ca^2+^ pumping and mode-switching by single LMCA1 pumps demonstrate the existence of a transprotein leakage pathway. The transprotein leakage lifetimes is ∼1-2 order of magnitude faster than that generated by ionomycin. Fits correspond to monoexponential functions. Errors correspond to the standard error of the fit using 95% confidence intervals.

**Figure S16.**
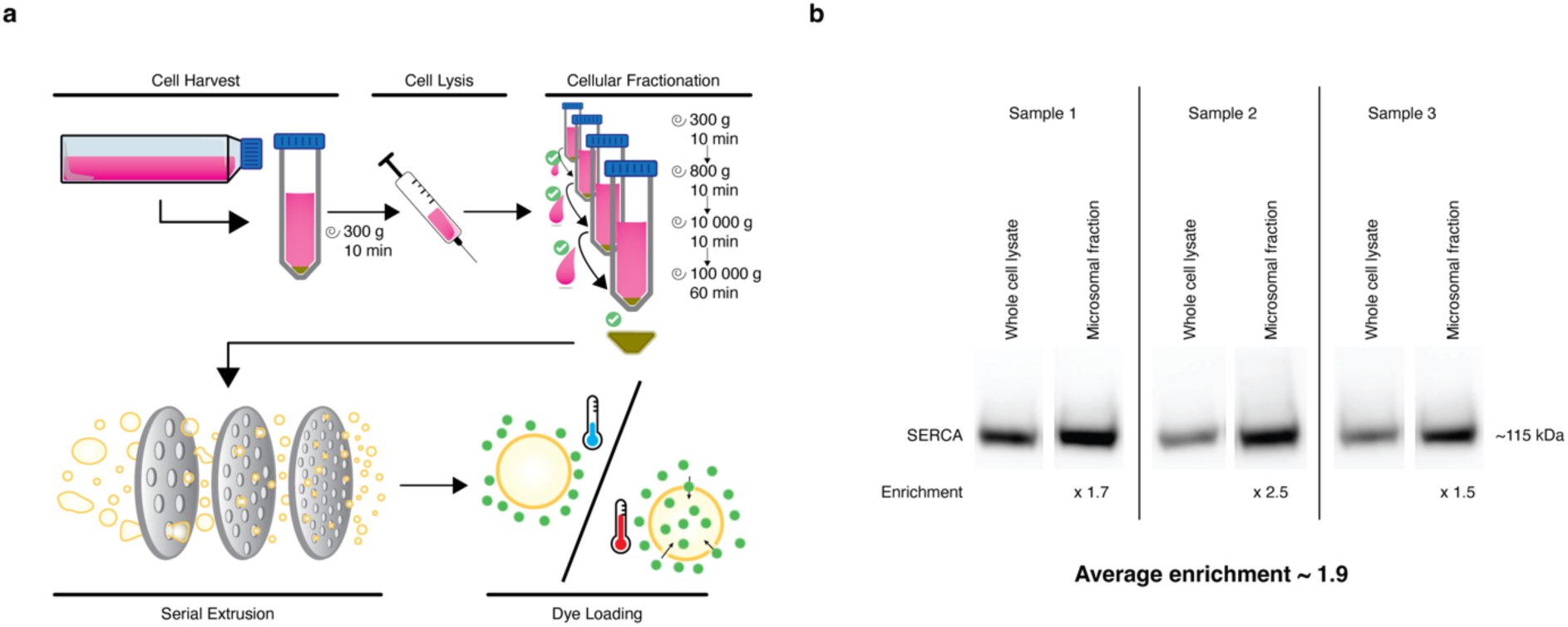
Method for generating native ER vesicles show enrichment of endogenously expressed hSERCA. **a**, HEK293 cells are use as source material for the generation of native vesicles. The cells are detached with a cell scraper, collected into a centrifuge tube and spun down. The pellet is resuspended in assay buffer and cells are lysed by repeated passage through a needle using a syringe. A classic subcellular fractionation protocol with several steps of differential ultracentrifugation is used to attain a microsome enriched fraction bearing SERCA molecules. This microsome fraction is extruded several rounds through consecutively smaller pore-size filters. Extruded native vesicles are then loaded with the calcium sensor Cal-520 by one round of freeze/thawing. Native vesicles are then immobilized on glass slides functionalized with BSA and biotin conjugated BSA in a 100:1 molar ratio, to which neutravidin is added as a linker between surface functionalized biotin-BSA and subsequently added biotin-PEG-cholesterol, which upon sample addition inserts into vesicle membranes and effectively tethers the vesicle to the surface. These surface anchored vesicles are now ready for activity measurements by TIRF microscopy. **b,** Western blots showing comparisons of SERCA specific bands between whole cell lysates and microsomal lysates of three separate isolates. Enrichment shows the fold increase of SERCA band densities in microsomal fractions as compared to that of whole cell lysates along with the average enrichment of three separate isolates.

**Figure S17.**
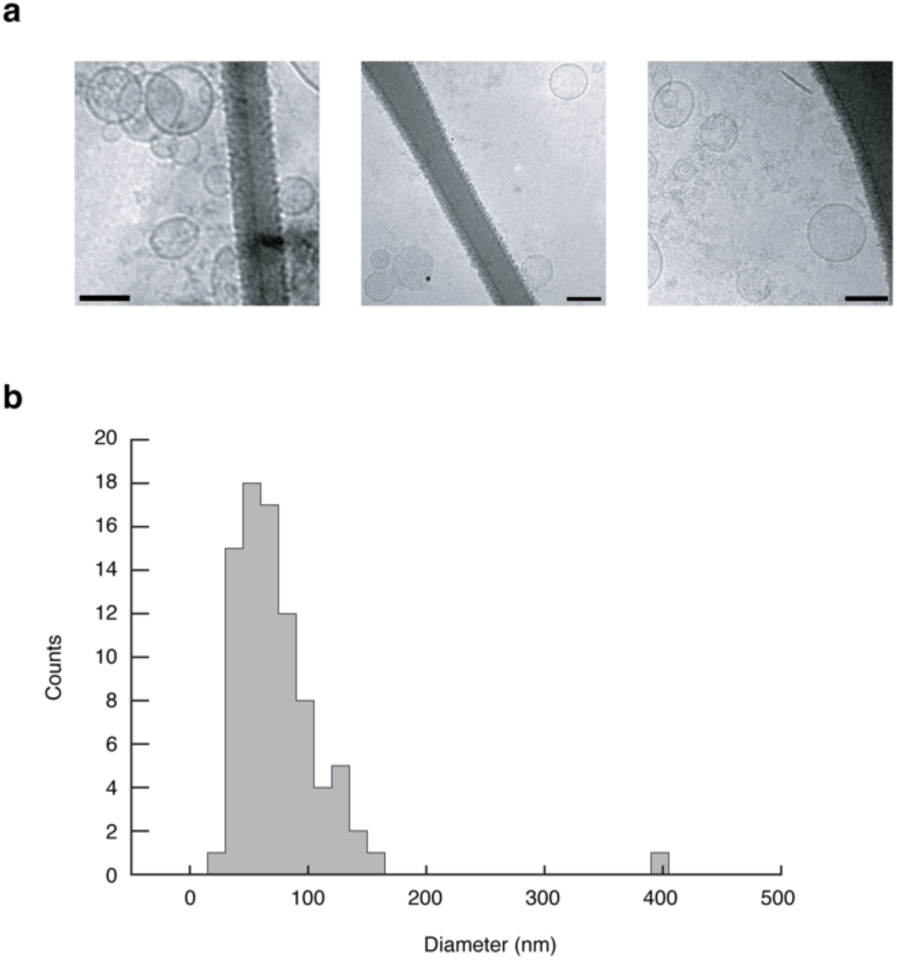
Size distribution and morphology of ER vesicles. **a**, Representative cryo-electron tomographs of native ER vesicles after extrusion. All scale bars correspond to 100 nm. **b,** Distribution of vesicle sizes after sequential extrusion through pores down to 100 nm demonstrates a mean vesicle diameter ∼70 nm and a monodisperse, homogeneous sample.

**Figure S18.**
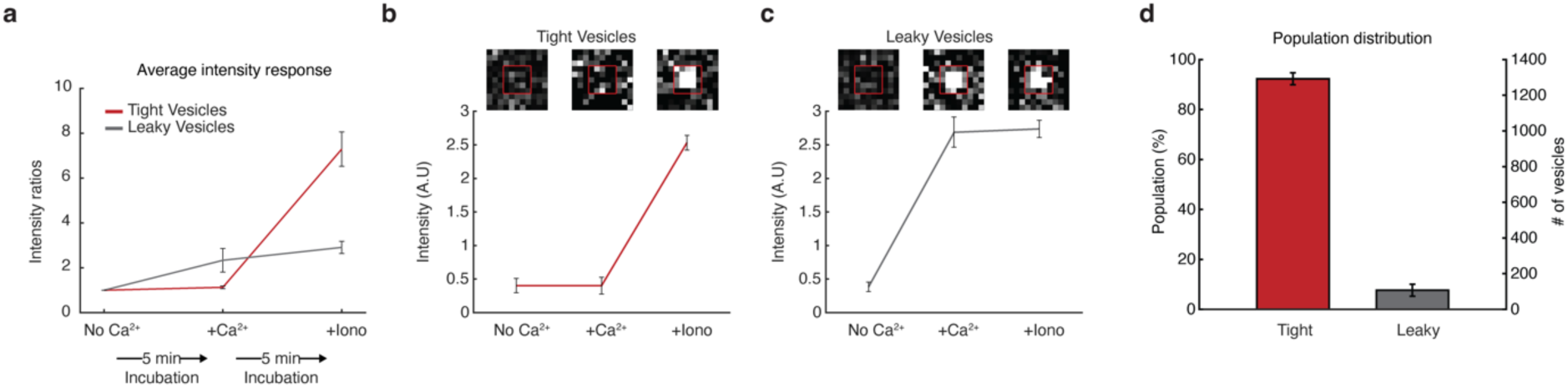
92% of ER vesicles are inherently impermeable to Ca^2+^. **a**, Ensemble average calculations of intensity ratios (± Ca^2+^) of single vesicles that are sequentially subjected and recorded (1) in the absence of calcium; (2) upon addition of 5 µM Ca^2+^; and (3) upon addition of 5 µM Ca^2+^ and 1 µM ionomycin. **b, c,** Representative examples of a tight (b) and a leaky (c) vesicle responding to Ca^2+^ and ionomycin. **d,** Percentage of sensor-loaded tight native vesicles constitute 92% of the identified vesicle population. Error bars correspond to s.d., *n* = 3.

**Figure S19.**
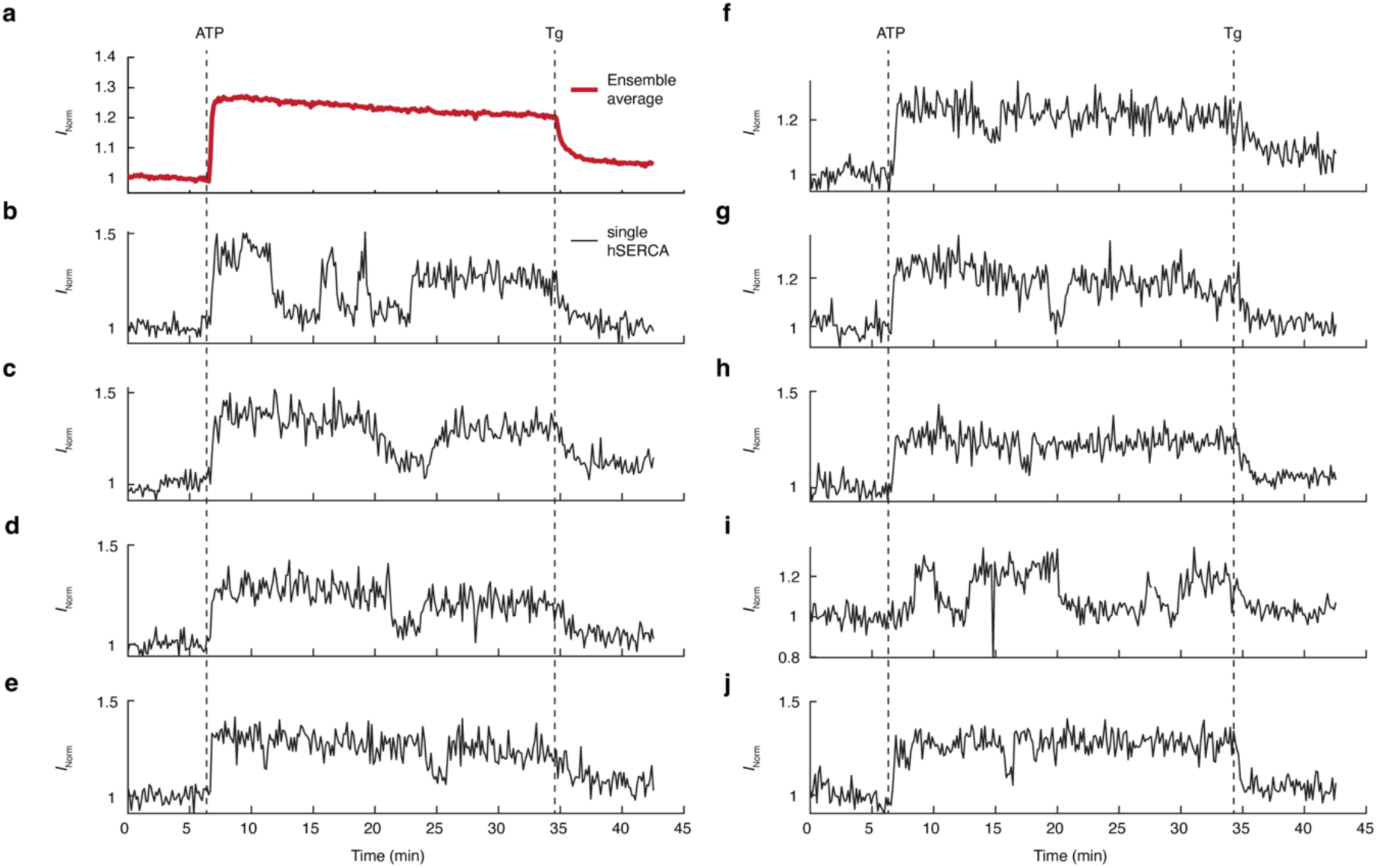
Ensemble average and single-molecule Ca^2+^ pumping kinetics in endogenous ER vesicles by hSERCA demonstrate mode-switching and are inhibited by Thapsigargin. **a**, Ensemble average of 208 single-vesicle Ca^2+^ pumping kinetics show that activity is initiated by ATP and is blocked upon addition of the SERCA-specific inhibitor Thapsigargin (Tg). **b-j,** Representative single-molecule kinetics show stochastic switching between active and inactive modes. These kinetic traces were not corrected for photobleaching.

**Fig. S20.**
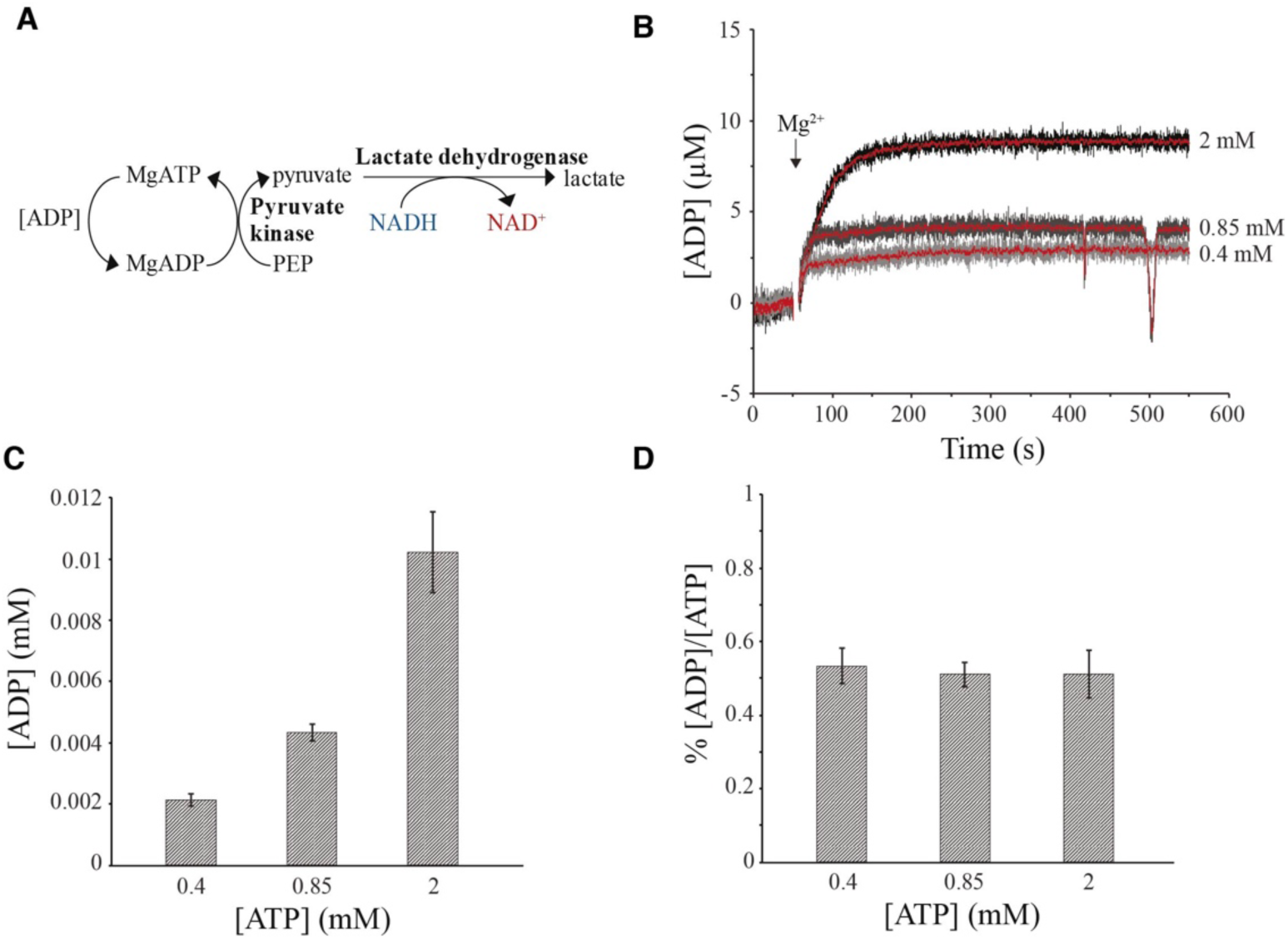
Determination of ADP contamination and ATP autohydrolysis. A) Schematic representation of a NADH coupled ATPase assay. B) Three ATP blanks containing 0.4, 0.85, and 2 mM ATP were assayed for ADP content using a NADH coupled ATPase assay. The graph shows representative background-subtracted traces from a continuous monitoring of the amount of ADP regenerated into ATP over 500 seconds. The ADP regeneration is triggered by addition of Mg^2+^. The red traces are moving averages using a period of 10. The initial increase in regenerated ADP originates from the conversion of ADP impurities in the ATP stock into ATP. After about 150 s, any further increase in the amount of regenerated ADP will be a result of ATP autohydrolysis during the measurement. As seen the slopes at the end of the traces are practically identical for the three different ATP concentrations and close to zero, excluding any detectable ATP autohydrolysis on the time scale of these measurements. C) The total amount of regenerated ADP after 500 seconds measured from the similar curves that are represented in B). Data represent mean ± SD (n=3), which is interpreted as the total amount of ADP contamination of the ATP stock. When calculated as percentage of ADP/ATP, values of 0.53, 0.51, and 0.51% are obtained for 0.4, 0.85, and 2 mM ATP respectively, as shown in D).

---

**Table S1.** Simulation parameters.

| Variable | Symbol | Value | Unit |
| --- | --- | --- | --- |
| Pumping rate<br>*Simulations*<br>*Experiments* | I | Variable<br>Model output | Ca<sup>2+</sup>/s |
| Membrane permeability<br>*Simulations*<br>*Experiments* | P<sub>mem</sub> | Variable<br>Model output | cm/s |
| Vesicle radius<br>*Simulations*<br>*Experiments* | r | 30, 100, 1000<br>Experiment | nm |
| Transprotein permeability | P<sub>LMCA1</sub> | Model output | cm/s |
| Cal-520 concentration | [Cal-520] | 2 | mM |
| Cal-520 dissociation constant | K<sub>d</sub> | 4.4 | μM |
| Extraluminal potassium concentration | [K<sup>+</sup>]<sub>o</sub> | 200 | mM |
| Luminal potassium concentration | [K<sup>+</sup>]<sub>i</sub> | 200 | mM |
| Bilayer capacitance | C<sub>0</sub> | 1 | F/cm<sup>2</sup> |
| Potassium permeability | P<sub>K</sub> | 10<sup>-7</sup> | cm/s |
| Donnan particles | B | 200 | mM |

